# Genetic differences in the aryl hydrocarbon receptor and CYP1A2 affect susceptibility to developmental polychlorinated biphenyl exposure in mice: Relevance to studies of human neurological disorders

**DOI:** 10.1101/194472

**Authors:** Kelsey Klinefelter, Molly Kromme Hooven, Chloe Bates, Breann T. Colter, Alexandra Dailey, Smitha Krishnan Infante, Izabela Kania-Korwel, Hans-Joachim Lehmler, Alejandro López-Juárez, Clare Pickering Ludwig, Christine Perdan Curran

## Abstract

Polychlorinated biphenyls (PCBs) are persistent organic pollutants that remain a human health concern with the discovery of new sources of contamination and ongoing bioaccumulation and biomagnification. Children exposed during early brain development are at highest risk of neurological deficits, but there is some evidence that high PCB exposure in adults increases the risk of Parkinson’s disease. Our previous studies found allelic differences in the aryl hydrocarbon receptor and cytochrome P450 1A2 (CYP1A2) affect susceptibility to developmental PCB exposure, resulting in cognitive deficits and motor dysfunction. High-affinity *Ahr*^*b*^*Cyp1a2(-/-)* mice were most susceptible compared with poor-affinity *Ahr*^*d*^*Cyp1a2(-/-)* and wild type *Ahr*^*b*^*Cyp1a2(+/+)* mice. Our follow-up studies assessed biochemical, histological and gene expression changes to identify the brain regions and pathways affected. We also measured PCB and metabolite levels in multiple tissues to determine if genotype altered toxicokinetics. We found evidence of AHR-mediated immune suppression with reduced thymus and spleen weights and significantly reduced thyroxine at P14. In the brain, the greatest changes were seen in the cerebellum where a foliation defect was over-represented in *Cyp1a2(-/-)* mice. In contrast, we found no difference in tyrosine hydroxylase immuno-staining in the striatum. Gene expression patterns varied across the three genotypes, but there was clear evidence of AHR activation. Distribution of parent PCB congeners also varied by genotype with strikingly high levels of PCB 77 in poor-affinity *Ahr*^*d*^*Cyp1a2(-/-)* while *Ahr*^*b*^*Cyp1a2(+/+)* mice effectively sequestered coplanar PCBs in the liver. Together, our data suggest that the AHR pathway plays a role in developmental PCB neurotoxicity, but we found little evidence that developmental exposure is a risk factor for Parkinson’s disease.

## Introduction

Polychlorinated biphenyls are persistent organic pollutants linked to numerous adverse health effects in humans, including immune suppression, endocrine disruption, and neurological deficits (ATSDR 2015). Despite numerous bans on commercial production, PCBs remain ubiquitous in the environment and the food supply (Gomara et al. 2005; Langer et al. 2007; Malisch and Kotz 2014; Marek et al. 2017). New PCB sources have recently been identified, including their inadvertent production during the manufacture of paint pigments (Hu & Hornbuckle 2010) and the discovery of PCBs in and around many U.S. schools (Marek et al. 2017). Because PCBs are lipophilic and can bioaccumulate and biomagnify, the risk to pregnant women and their offspring is expected to remain high for years to come (Quinn et al. 2011; Bányiová et al. 2017).

The evidence for PCB-induced motor dysfunction comes from both human and animal studies. Stewart et al. (2000), Vreugdenhil et al. (2002), and Boucher et al. (2016) all reported motor deficits in children exposed to high levels of PCBs. Hydroxylated PCB metabolites are also linked with motor dysfunction (Berghuis et al. 2013; Haijima et al. 2017). Steenland et al. (2006) reported a higher risk of Parkinson’s disease (PD) in women with high occupational exposure. Similarly, Petersen et al. (2008) reported a higher risk of PD in adults with high consumption of contaminated whale meat and blubber. The risk of PD is consistent with animal studies showing dopamine depletion after PCB exposure (Seegal et al. 1994, 1997, 2005).

However, PCBs are a diverse group of chemicals with 209 possible congeners and multiple targets in the body. PCBs are well known to deplete thyroid hormone levels (Curran et al. 2011a; Giera et al. 2011), and supplemental thyroxine rescues PCB-induced motor deficits (Goldey and Crofton 1998). Interestingly, CYP1A1 can metabolize some PCB congeners into hydroxylated metabolites that disrupt thyroid hormone signaling (Gauger et al. 2007; Wadzinski et al. 2014). Since thyroid hormone is essential for normal cerebellar development (Zoeller and Rovett 2004), it is possible that at least some of the observed motor deficits could be caused by cerebellar dysfunction and not deficits in the nigrostriatal pathways affected in Parkinson’s disease.

Our previous work demonstrated that allelic differences in the aryl hydrocarbon receptor (AHR) and cytochrome P450 1A2 (CYP1A2) alter susceptibility to developmental PCB neurotoxicity, leading to greater impairments in learning and memory (Curran et al. 2011b, 2012) and impaired motor function in *Ahr*^*b*^*Cyp1a2(-/-)* knockout mice (Stegman et al. 2014; Colter et al. 2017 submitted). Both of these genes are highly polymorphic in the human population (Nebert & Dalton 2006), so our model mimics human genetic variation. The focus of this project was to further characterize genetic susceptibility to PCBs and to explore mechanisms of neurotoxicity by examining changes in thyroid hormone signaling and gene expression in brain regions required for normal motor function: the dorsal striatum, cerebellum and cortex.

## Materials and Methods

### Animals

We used C57BL/6J mice (The Jackson Laboratory, Bar Harbor ME) with the *Ahr*^*b*^*Cyp1a2(+/+)* genotype and two lines of knockout mice which have been back-crossed at least 8 generations onto the B6 background: high-affinity *Ahr*^*b*^*Cyp1a2(-/-)* and poor-affinity *Ahr*^*d*^*Cyp1a2(-/-)* mice. All mice were housed in polysulfone shoebox cages with corncob bedding and one 5 cm^2^ nestlet provided weekly for enrichment. Food (LabDiet 5015) and water were provided *ad libitum* with a 12h:12h light dark cycle under temperature and humidity-controlled conditions. A maximum of one male and one female per litter was used in each test (N ≥ 6 litters per group for all tests).

### Breeding

Nulliparous females between 2.5 and 4 months of age were mated with males of the same genotype in a 4-day breeding cycle. The morning a vaginal plug was found, the female was separated from the male and housed individually for delivery.

### Treatments

An environmentally relevant mixture of PCBs (coplanar PCBs 77, 126, 169 and noncoplanar PCBs 105, 118, 138, 153 and 180, ULTRA Scientific, North Kingstown RI) were dissolved in corn oil, and dams were dosed daily by providing a small piece of Froot Loop® cereal containing the PCB solution or plain corn oil for control animals. Additional information regarding the dosing solutions is provided as Supplemental Material (Table S1). Dosing began the afternoon of the day a vaginal plug was found and continued through postnatal day 25 (P25) with animals randomly assigned to each group.

### Tissues

To avoid confounding by circadian rhythms, all tissues were collected during the same 3h time period (15:00-18:00h) at P14, P25, P30 and P120. Blood was collected by cardiac puncture for PCB quantification. Trunk blood was collected for thyroid hormone measurements, centrifuged at 1500G for 5 min at 4°C, and the plasma fraction stored at -80°C until analysis. Liver, inguinal fat pad, cerebellum, and cortex were harvested, snap frozen and stored at -80°C until analysis. Spleen and thymus were dissected out in P14 mice and wet weights recorded.

### Thyroid hormone measurements

Total thyroxine (T4) levels were measured using a standard ELISA kit (Alpco Diagnostics, Salem NH) following the manufacturer’s protocol. We measured T4 levels at P14 and in P120 littermates used in motor function tests.

### Tissue processing for immunohistochemistry

Mice were euthanized with CO2 and perfused transcardially with 1X PBS followed by 4% paraformaldehyde in phosphate buffer. Brains were removed and cryoprotected in 30% sucrose in 0.1 M PBS for coronal sectioning on a cryostat (striatum) or sagittal sectioning on a vibratome (cerebellum). Brains were collected at P25 and at P120 from littermates used for motor function testing.

### Immunohistochemistry for tyrosine hydroxylase in striatum

Six striatal sections per animal were selected for immunohistochemistry. To enhance antigen retrieval, we soaked tissue in sodium citrate buffer at 80° C for 30 min followed by washes in 0.1 M PBS. Next, tissues were incubated in 0.3% H_2_O_2_ (Sigma) in 0.1M PBS for 30 min at room temperature with shaking. The solution was replaced with normal goat serum/Triton 100X blocking solution and incubated for 2 h at room temperature with shaking. The tissue was then incubated in the primary antibody (Abcam Cat# ab112, RRID:AB_297840) at a 1:5000 dilution in blocking solution at 4° C for 48 h with shaking. After washes to remove the primary antibody, the slices were incubated for 2 h in the secondary antibody (anti-rabbit IgG biotinylated antibody from Vector ABC Elite kit) at room temperature with shaking. The slices were then incubated in HRP-ABC complex (Vector ABC Elite) for 1 h at room temperature with shaking followed by staining with diaminobenzidine (Vector DAB kit) and incubation for 10 min at room temperature. Following a final 15-min rinse sequence with 0.1M PBS, the sections were stored in 0.1M PBS with 0.05% sodium azide until mounting.

### Optical density analysis

Slides were viewed on an Olympus microscope with a Motic Camera attachment at 10X magnification and associated software (Version 2.0). Slides were photographed in grayscale for importing into ImageJ. All images were calibrated using the standard ImageJ step tablet to ensure uniformity. Background was subtracted using the rolling ball radius function. A grid was overlaid on the image and the line closest to the center of the striatum as possible was selected. This area was divided evenly into left and right dorsal and ventral quadrants. We used a random number generator to select six squares from each hemisphere for analysis (Fig. 1). The optical density of each square was measured, and the mean O.D. for each hemisphere was calculated for statistical analysis following the methods of (González-Franco et al. 2017).

**Fig. 1.**
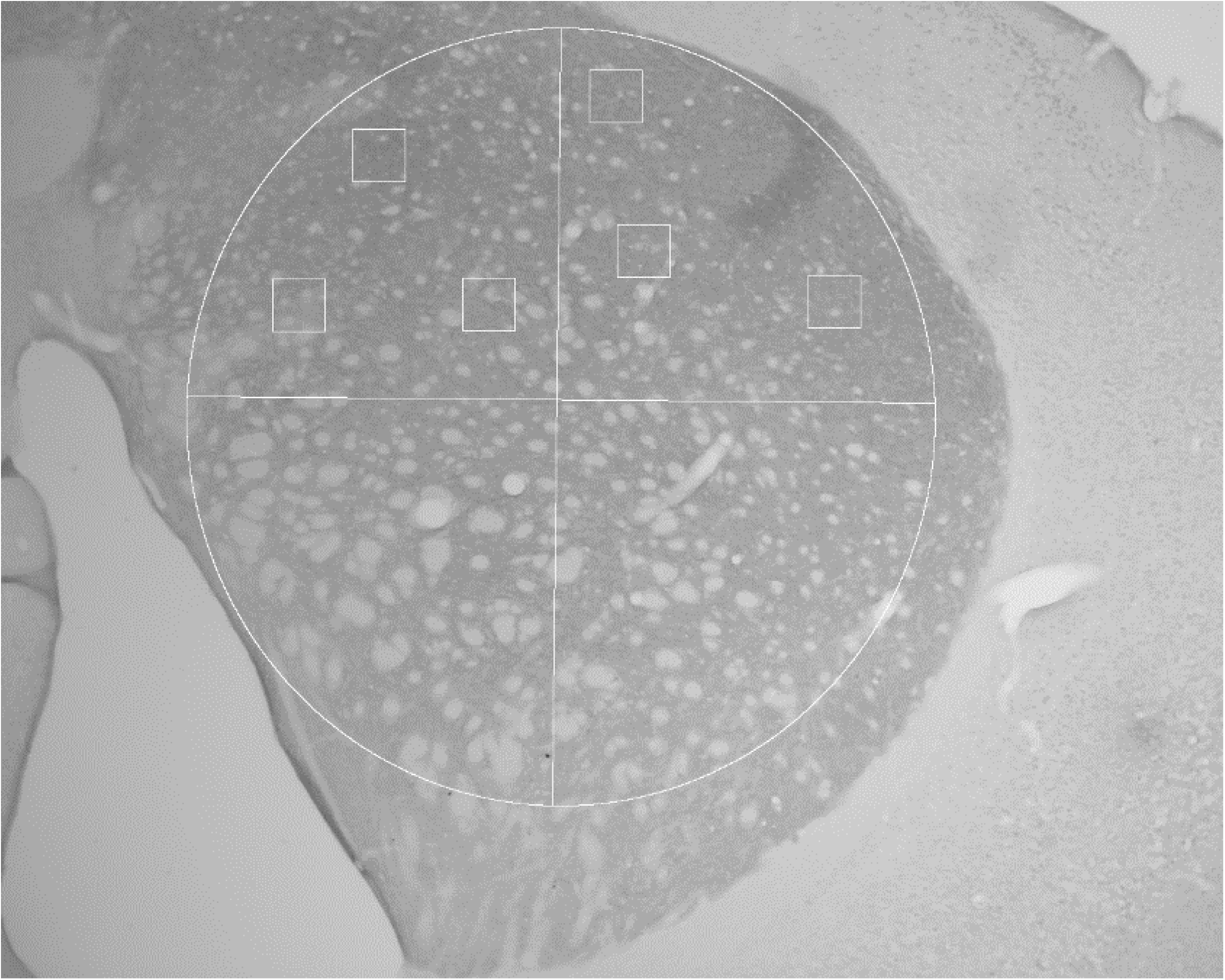
Sampling scheme for optical density measurements in the dorsal striatum. Modified from González-Franco et al. 2017.

### Immunohistochemistry and image analysis of cerebellum

Vibratome floating sections were blocked in PBS 1X/Triton 100X 0.3%/FBS and processed for immunodetection of proteins using specific antibodies for GFAP (Sigma-Aldrich; G-3893), Blbp (Millipore; ABN14) Parvalbumin (Millipore; AB15738), Calbindin (Sigma-Aldrich; C8666), and NeuN (Millipore; MAB377). The appropriate fluorophore-conjugated secondary antibodies (Alexa 488, Alexa 568, Alexa 647; Invitrogen) were used to detect the antigen-antibody complexes. Nuclei were stained with DAPI, and sections were mounted on slides using Fluoromount G (Thermo Fisher). Using the appropriate laser excitation wavelength (405, 488 or 561 nm), images were captured on a Nikon C2 confocal microscope. ImageJ (NIH) software was used to calculate the perimeter and area of the granule cell layer and to compare the number of Purkinje cells across different folia. Images were assessed by investigators blinded to genotype and treatment groups.

### Real-time quantitative PCR arrays

Tissues were homogenized in RNAzol (Molecular Research Center, Cincinnati, OH) with 50-100 mg tissue per ml RNAzol using a Dounce homogenizer for cortex and cerebellum and a Polytron-type homogenizer for liver. RNA was extracted using the manufacturer’s protocol, and the RNA pellets were re-suspended in 50 μl of nuclease-free water for brain and in 75 μl for liver. RNA concentration was quantified on a Nanodrop (Thermo Fisher), and 500 ng total RNA was used for the cDNA reaction using the RT^2^ First Strand Kit (Qiagen).

The cDNA was analyzed using an RT^2^ Custom PCR Array (Qiagen) with primers for 39 candidate genes, 5 housekeeping genes, and negative and positive controls. Genes were selected after a literature search for genes related to activation of the aryl hydrocarbon receptor, cerebellar development, oxidative stress, neuroinflammation, and other pathways implicated in PCB toxicity and endocrine disruption (ryanodine receptor/calcium homeostasis, estrogen and thyroid hormone pathways). Two samples were run on each 96-well plate in an Applied Biosystems 7300 Real-time PCR system using the following program: 10 min at 95°C, 40 cycles of 15 s at 95°C followed by 1 min at 60°C. The RT^2^ PCR Array Data Analysis version 3.5 was utilized to compare fold-change in three genotypes of PCB-treated mice with corn oil-treated wild-type mice using the Delta-Delta Ct method.

### Chemicals for PCB quantification

Analytical standards of 2,3,5,6-tetrachlorobiphenyl (PCB 65, 99.4%); 2,3,3′,4,4′-pentachlorobiphenyl (PCB 105, > 99.9%); 2,2′,3,4,4′,5′-hexachlorobiphenyl (PCB 138, > 99.9%); 2,3,4,4′,5,6-hexachlorobiphenyl (PCB 166, 99%); 3,3′,4,4′,5,5′-hexachlorobiphenyl (PCB 169, 99%); 2,2′,3,4,4′,5,5′-heptachlorobiphenyl (PCB 180, 99.2%), 2,2′,3,4,4′,5,6,6′-octachlorobiphenyl (PCB 204, 99.9%) and 2′,3,3′,4′,5,5′-hexachlorobiphenyl-4-ol (4-159, > 99.9%) were purchased from AccuStandard (New Haven, CT). Standards of 3,3′,4,4′-tetrachlorobiphenyl (PCB 77); 2,3′,4,4′,5-pentachlorobiphenyl (PCB 118); 3,3′,4,4′,5-pentachlorobiphenyl (PCB 126) and 2,2′,4,4′,5,5′-hexachlorobiphenyl (PCB 153) were synthesized as described previously (Kania-Korwel et al. 2004, Lehmler and Robertson 2001, Schramm et al. 1985). Methylated standards of OH-PCBs, including 2,3,3′,4′,5-pentachloro-4-methoxybiphenyl (4-107, >98%); 2,2′,3′,4,4′5-hexachloro-3-methoxybiphenyl (3′-138, >98%); 2,2′,3,4′,5,5′-hexachloro-4-methoxybiphenyl (4-146, >98%) and 2,2′,3,4′,5,5′,6-heptachloro-4-methoxybiphenyl (4-187, >98%), were purchased from Wellington Laboratories (Guelph, ON), Canada. Hydroxylated metabolite standards of PCB 77, including 3,3′,4′,5-tetrachloro-biphenyl-4-ol (4′-79); 3′,4,4′,5-tetrachlorobiphenyl-3-ol (5-77); and 3′,4,4′,5-tetrachlorobiphenyl-2-ol (6-77) were prepared as described previously (van den Hurk et al. 2002). Standards synthesized in our laboratory were >99% pure by GC analysis. The nomenclature of metabolites follows an abbreviated nomenclature proposed by Maervoet et al. (2004).

### Extraction of tissues

Adipose (0.0606 – 0.3107 g), blood (0.2446 – 0.5884 g), cerebellum (0.0319 – 0.0573 g), cortex (0.1215 – 0.2468 g) and liver samples (0.2402 – 0.2468 g) were spiked with surrogate standards (PCBs 65 and 166, 4-159) and extracted with hexane-acetone (1:1 v/v) using an Accelerated Solvent Extractor (Dionex, Sunnyvale, CA) as described previously (Kania-Korwel et al. 2007). Sample blanks containing only Florisil and diatomaceous earth as well as an ongoing precision and recovery (OPR) standard (i.e, sample blanks spiked with all analytes) were extracted in parallel with each set of samples. Extracts were concentrated and derivatized with diazomethane to transform OH-PCBs into methoxylated compounds for gas chromatographic analysis (Kania-Korwel et al. 2008). Afterwards, samples underwent sulfur clean-up, followed by sulfuric acid treatment as reported previously (Kania-Korwel et al. 2005).

### Gas chromatography analysis

Levels of PCBs and OH-PCB metabolites in each extract were quantified using an Agilent 6890N or Agilent 7890A gas chromatograph equipped with a SPB-1 column 60 m length, 0.25 mm inner diameter, 0.25 μm film thickness; Supelco, St. Louis, MO) and a ^63^Ni μ-electron capture detector. The injector and detector temperatures were 280°C and 300°C, respectively. The following temperature program was used: 80°C for 1 min, 15°C/min increase to 260°C, 1°C/min increase to 274°C, 15°C/min increase to 300°C. PCBs 105+153 and 5-77+4-107 co-eluted on the SPB-1 column and are reported as sum of both co-eluting compounds. PCB and OH-PCB levels were calculated using PCB 204 as volume corrector and adjusted for tissue wet weights. The instrument and method detection limits for target analytes are summarized in Table S2.

### Quality assurance/quality control

The detector response was linear (R^2^ > 0.99) in the range from 1 to 500 ng for all analytes (R^2^ > 0.99). The recoveries of the surrogate standards PCB 65, PCB 166 and 4-159 were 88±11% (range: 53-112), 94±13% (range: 55 to 129) and 80±18% (range: 29 to 114), respectively. The concentrations of target analytes were corrected by the recoveries of the surrogate standards added to each sample. The recoveries from the OPR standard analyzed in parallel with each sample set ranged from 78 to 112%.

### Statistical analysis

Data were analyzed using SAS Proc Mixed Models Analysis of Variance with litter as the unit of analysis. When differences were found, we examined slice effects with a correction for multiple post-hoc analyses. For statistical analyses of PCB congeners and metabolites, values were set to the Limit of Detection for samples shown as Not Detected (ND). To compare the frequency of the cerebellar foliation deficit, we used a Chi Square analysis. Data are presented as least square means ± the standard error of the mean (SEM). For all analyses P < 0.05 was considered significant.

## Results

### Evidence of immune suppression

There was a main effect of PCB treatment and a significant gene x treatment interaction with PCB-treated P14 pups having significantly lower spleen (P < 0.01) and thymus (P < 0.001) wet weights. The greatest immune suppression was seen in PCB-treated *Ahr*^*b*^*Cyp1a2(-/-)* knockout mice (Figs. 2A-D).

**Figs. 2A-B Spleen wet weights.**
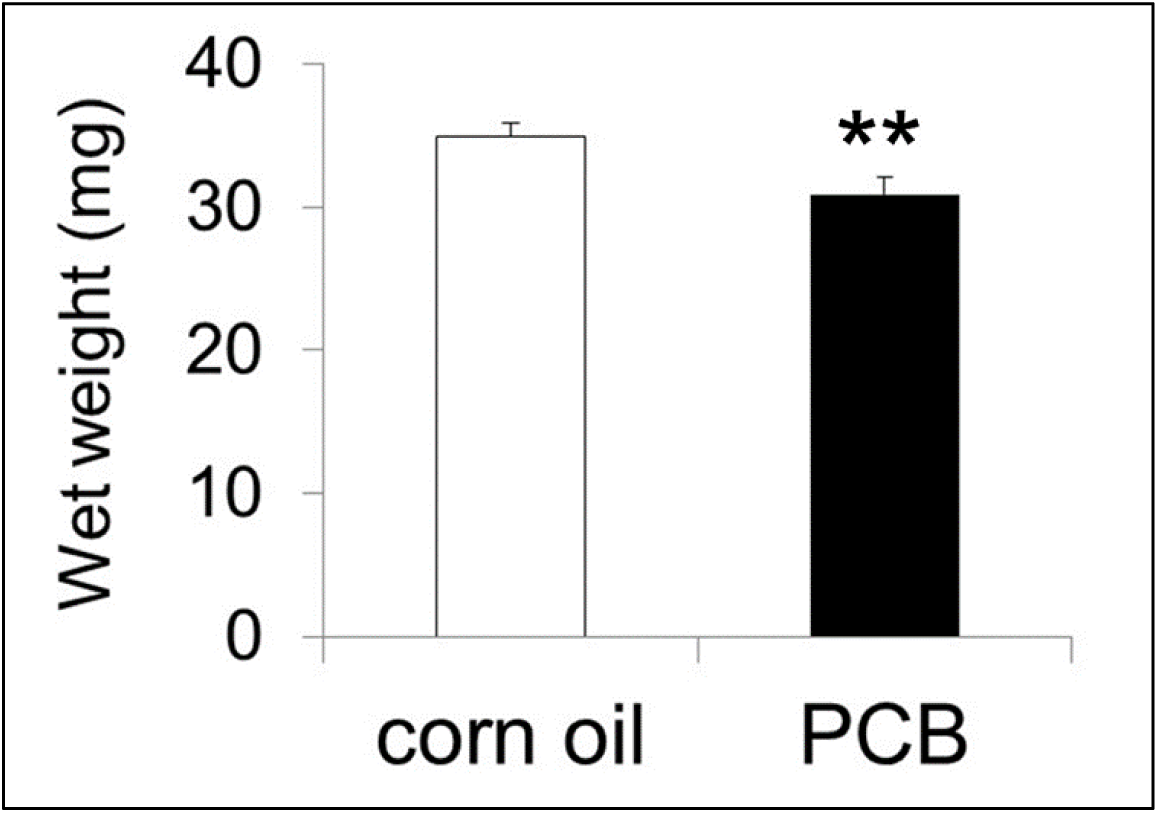
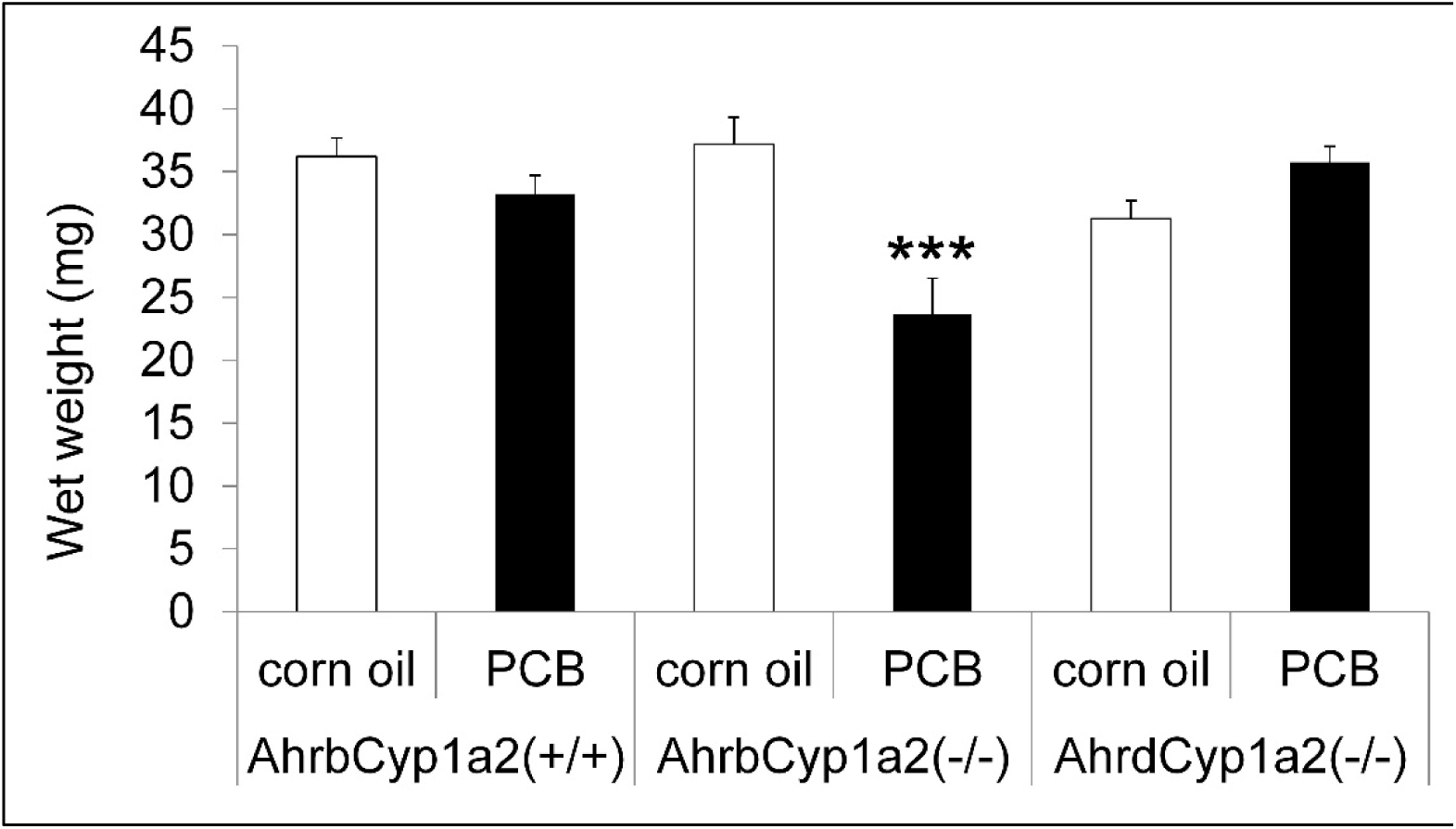
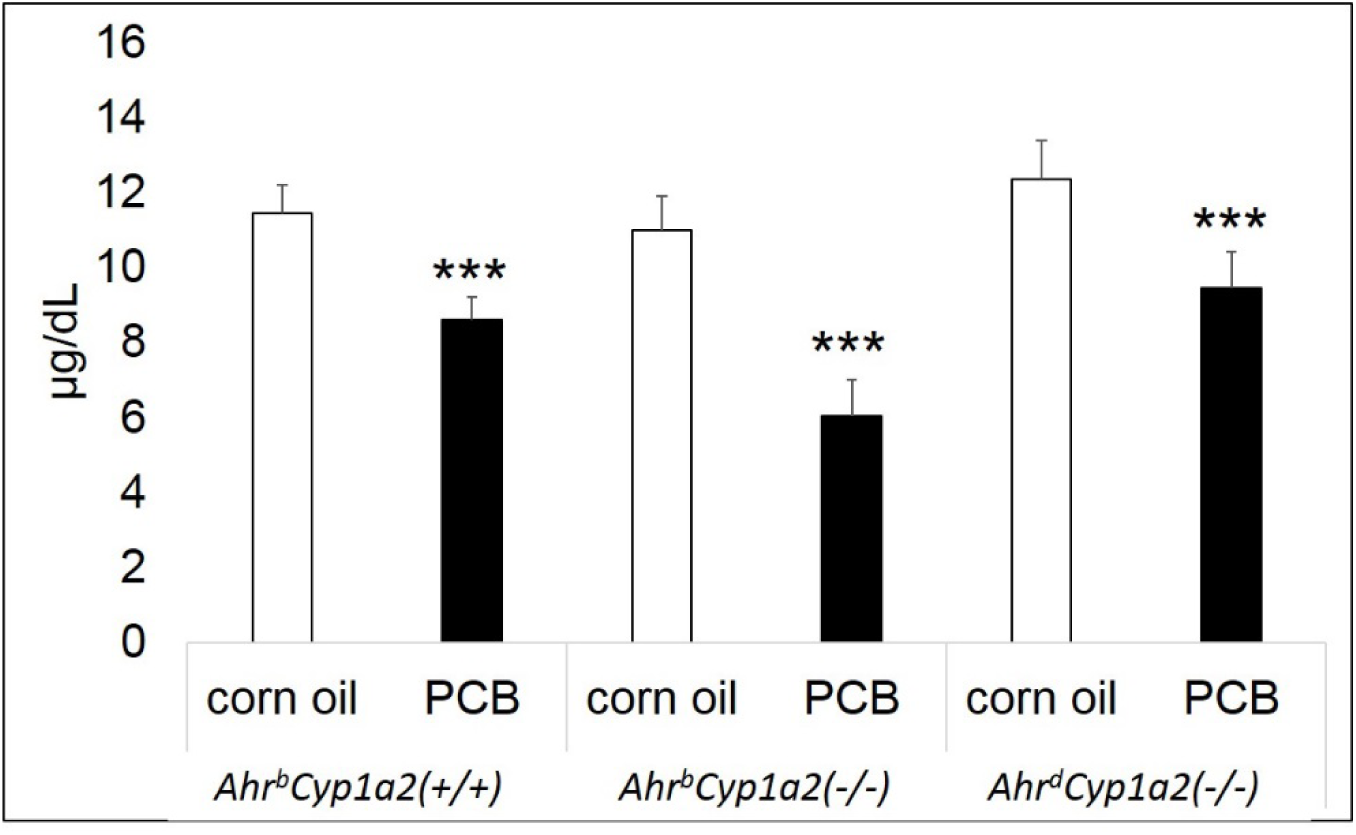
There was a main effect of treatment with PCB-exposed pups having significantly lower spleen weights at P14 compared to vehicle-treated controls. There was also a significant gene * treatment interaction with PCB-treated *Ahr*^*b*^*Cyp1a2(-/-)* knockout mice having significantly lower spleen weights than all other groups. ** P < 0.01, **A;* P < 0.001

**Figs. 2C-D Thymus wet weights.**
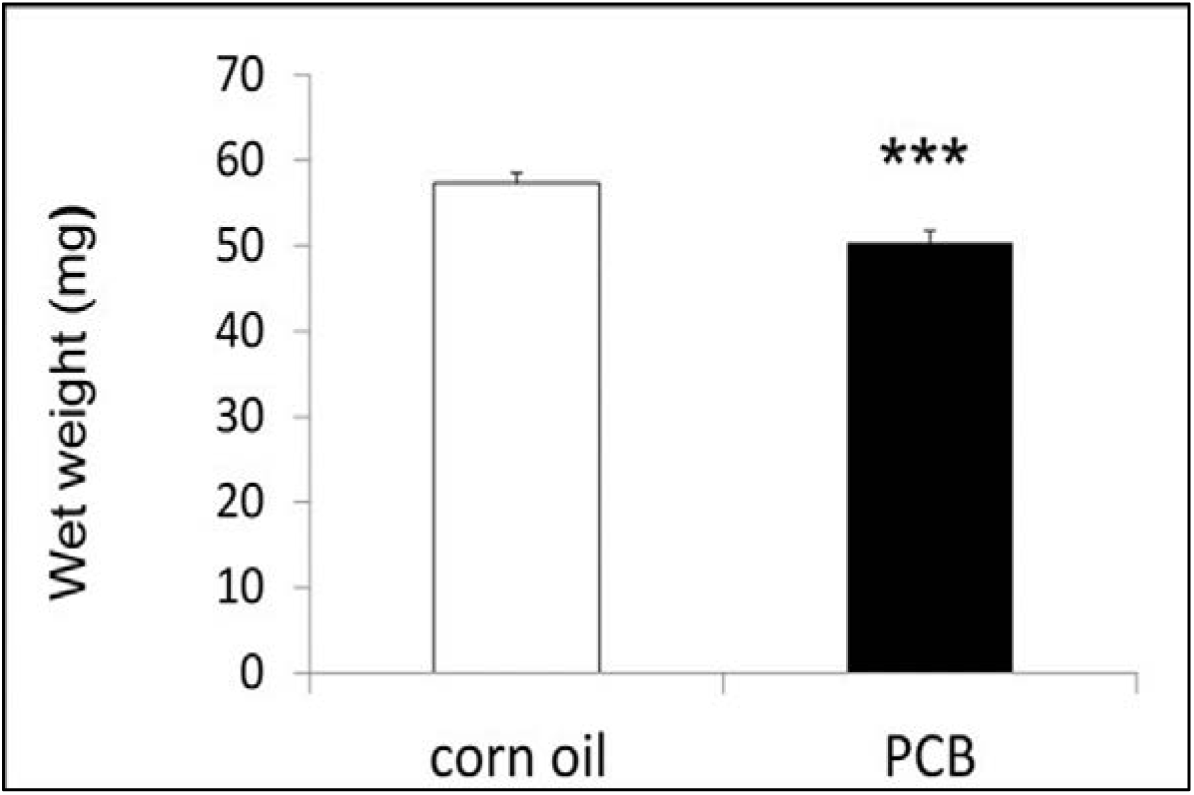
There was a main effect of treatment with PCB-exposed pups having significantly lower thymus weights at P14 compared to vehicle-treated controls. There was no gene * treatment interaction, although *Ahr*^*b*^*Cyp1a2(-/-)* knockout mice showed the greatest reduction compared to controls. **A;* P < 0.001

### Thyroid hormone levels

Total plasma thyroxine (T4) was significantly lower at P14 in all three genotypes of mice exposed to PCBs during gestation and lactation (P < 0.001); however, T4 levels returned to normal in adult mice where there were no significant differences (Figs. 3A-C).

**Figs. 3A-B Thyroxine levels in neonates.**
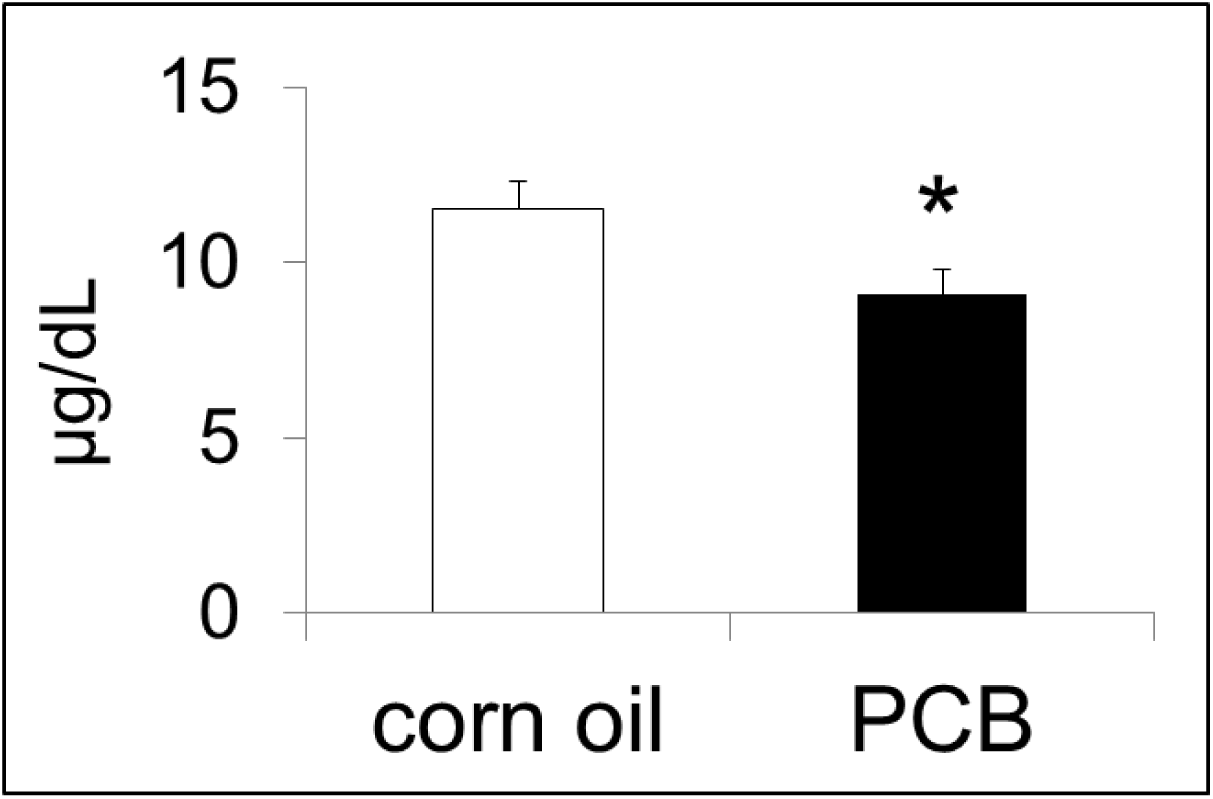
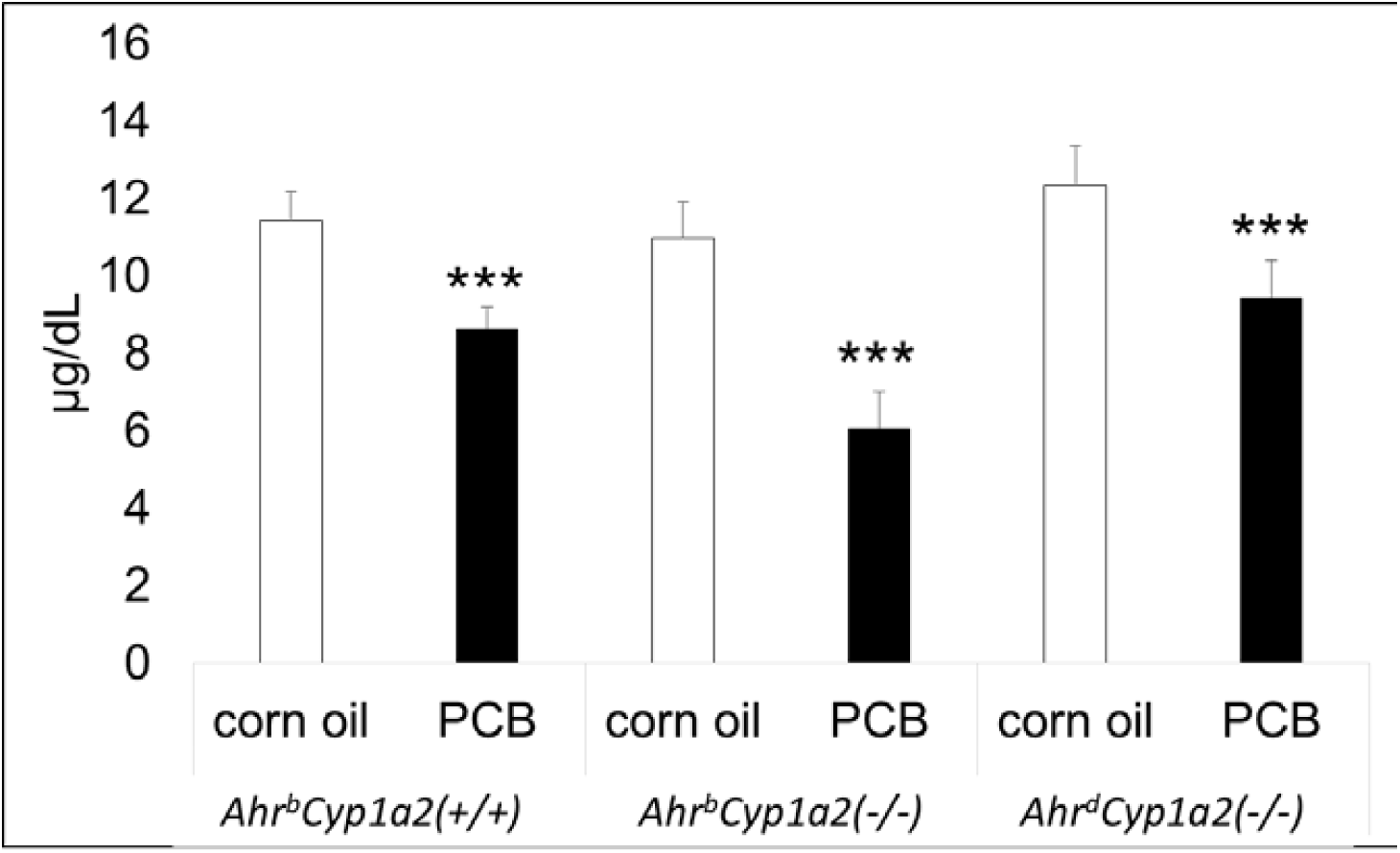
There was a main effect of treatment with PCB-exposed pups having significantly lower circulating free thyroxine at P14 compared to vehicle-treated controls. There was no gene * treatment interaction, although *Ahr*^*b*^*Cyp1a2(-/-)* knockout mice showed the greatest reduction compared to controls. **A;* P < 0.001

**Figs. 3A-B Thyroxine levels in adults.**
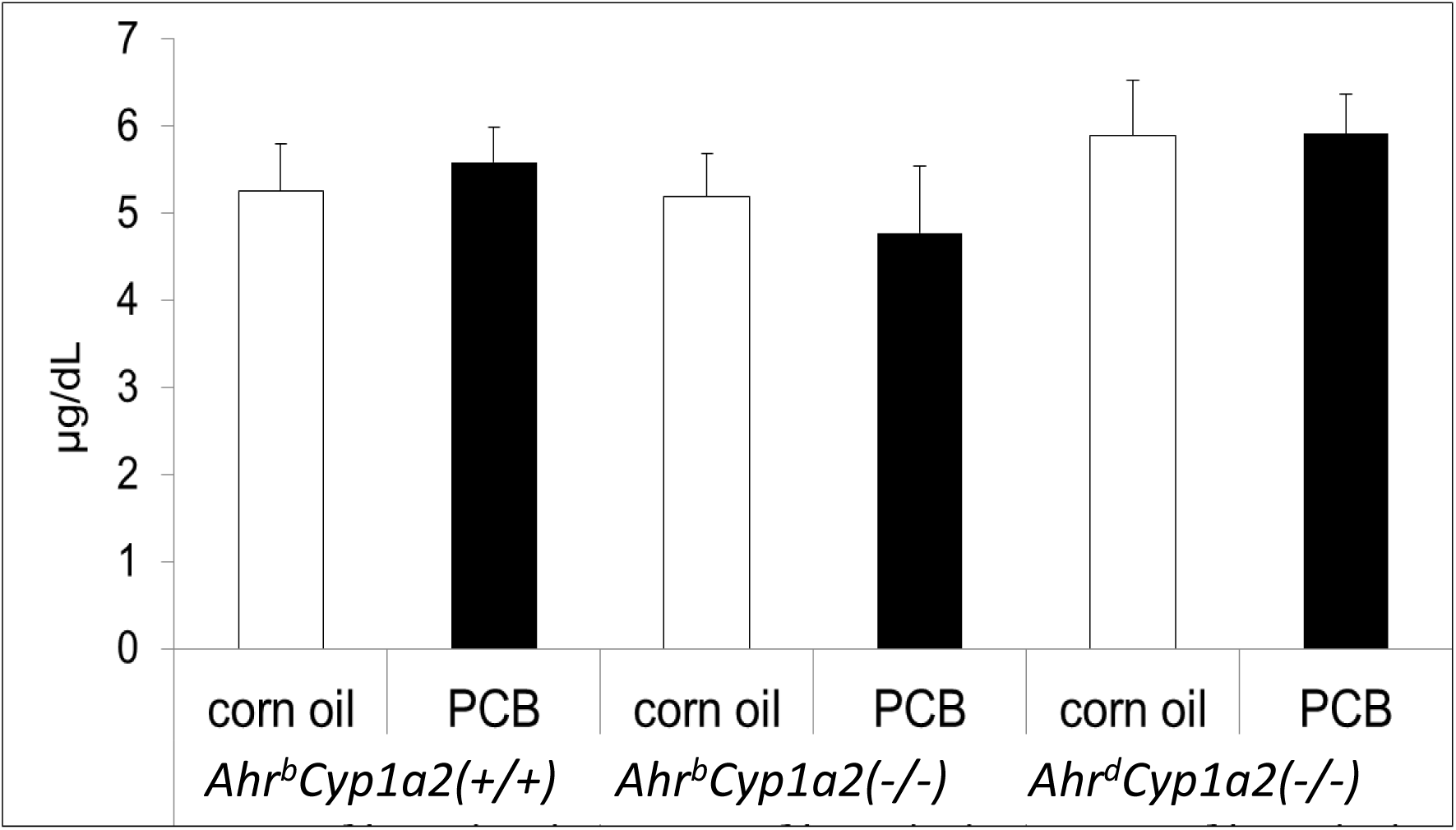
Free thyroxine levels returned to normal in adult mice tested at 4 months of age with no significant differences by genotype or treatment. P > 0.05

### Optical density measurements of striatum

We compared tyrosine hydroxylase staining in the striatum of PCB-treated mice from all three genotypes to values from corn oil-treated control B6 mice. There were no significant differences in tyrosine hydroxylase staining across these four groups (Fig. 4).

**Fig. 4 Optical density of dorsal striatum.**
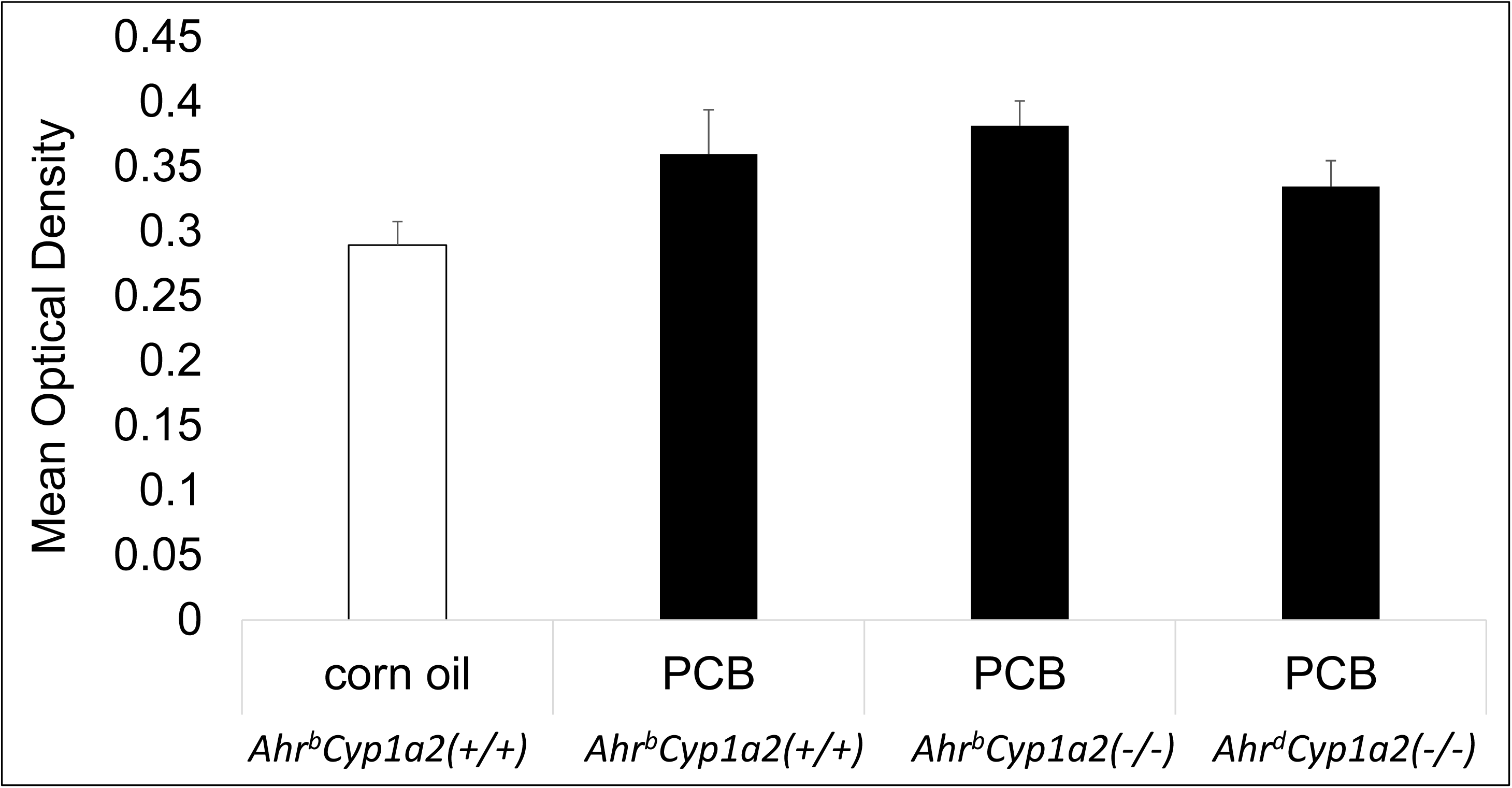
There were no significant differences in the mean optical density recorded following immuno-staining for tyrosine hydroxylase in the dorsal striatum. P > 0.05

### Histology of cerebellum

Decreases levels of thyroxine in PCB-treated mice (Figs. 3A-C) may affect biological processes where thyroid hormone is requested, such as cerebellum development. To test this possibility, we analyzed cerebellar structure and cellular composition in vehicle- and PCB-treated mice. We found no significant differences in the total area of the cerebellum at the level of the vermis, the area of the granule cell layer, the number of Purkinje cells, or the perimeter of the Purkinje cell layer in PCB-treated mice compared with their respective corn oil-treated controls (data not shown).

### Cerebellar foliation defect

A comparison of sagittal sections from control B6 mice and PCB-treated knockout mice revealed a foliation defect involving folia VIII and IX (Figs. 5A-B). This observation was followed up with a detailed analysis of cerebellar sections from all groups. We observed abnormal foliation patterns at a significantly higher frequency in PCB-treated mice (P < 0.01) with the highest percentage (67%) in PCB-treated *Ahr*^*b*^*Cyp1a2(-/-)* mice followed by 50% of PCB-treated *Ahr*^*d*^*Cyp1a2(-/-)* mice. In contrast, only 18% of corn oil-treated control mice showed the defect. In addition to aberrant lamination, we also detected invasion of Bergmann glial cells into the granule cell layer of PCB-treated knockout mice (Figs. 6A-F). These results suggest late cerebellar developmental defects in response to PCBs.

**Figs. 5A-B DAPI staining of cerebellum.**
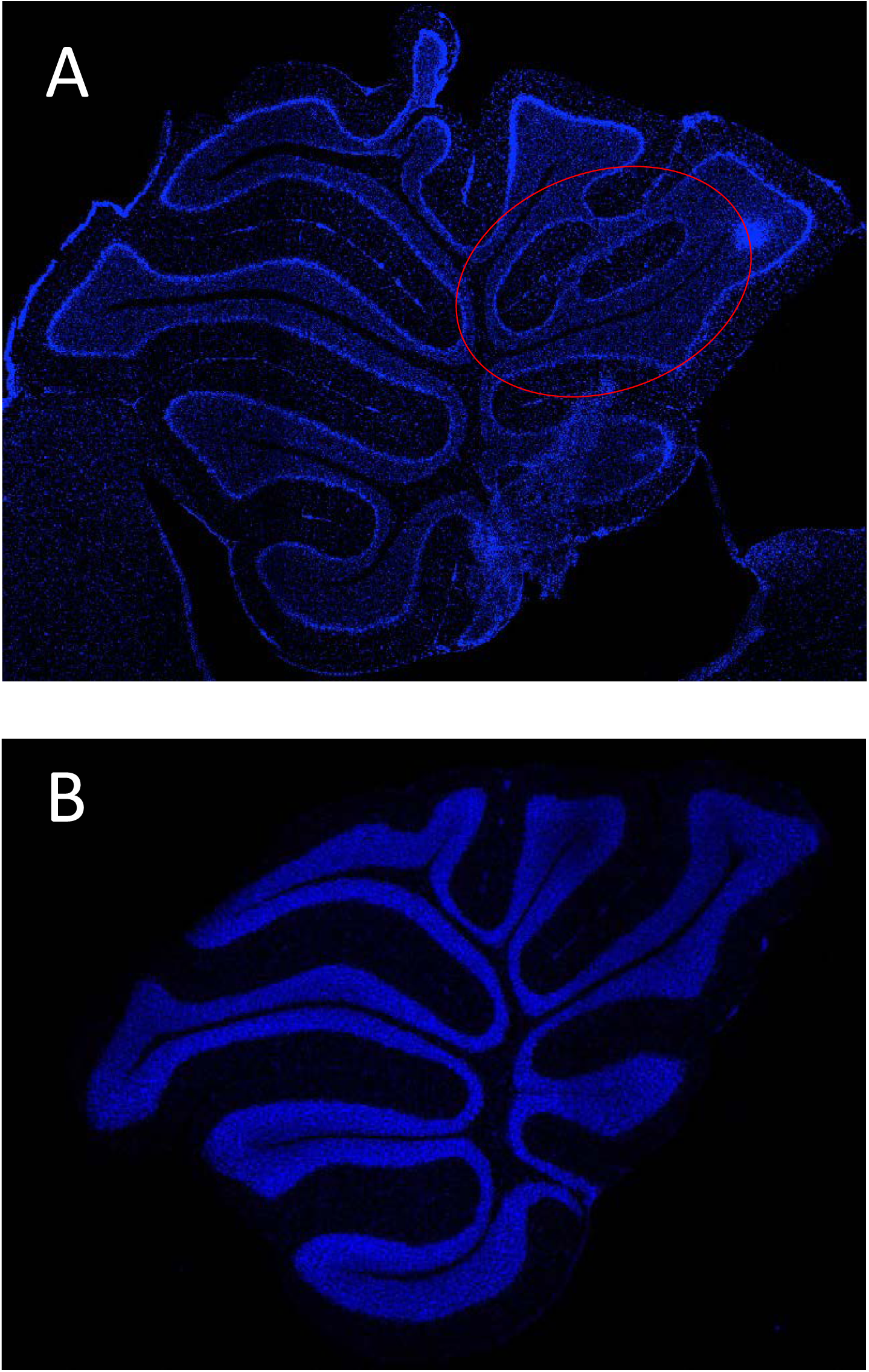
Representative sections from a PCB-treated *Ahr*^*b*^*Cyp1a2(-/-)* mouse (A) compared to a corn oil treated control *Ahr*^*b*^*Cyp1a2(+/+)* mouse (B). The region highlighted in red shows the foliation defect.

**Figs. 6A-F Defects in lamination pattern in folia VIII-IX.**
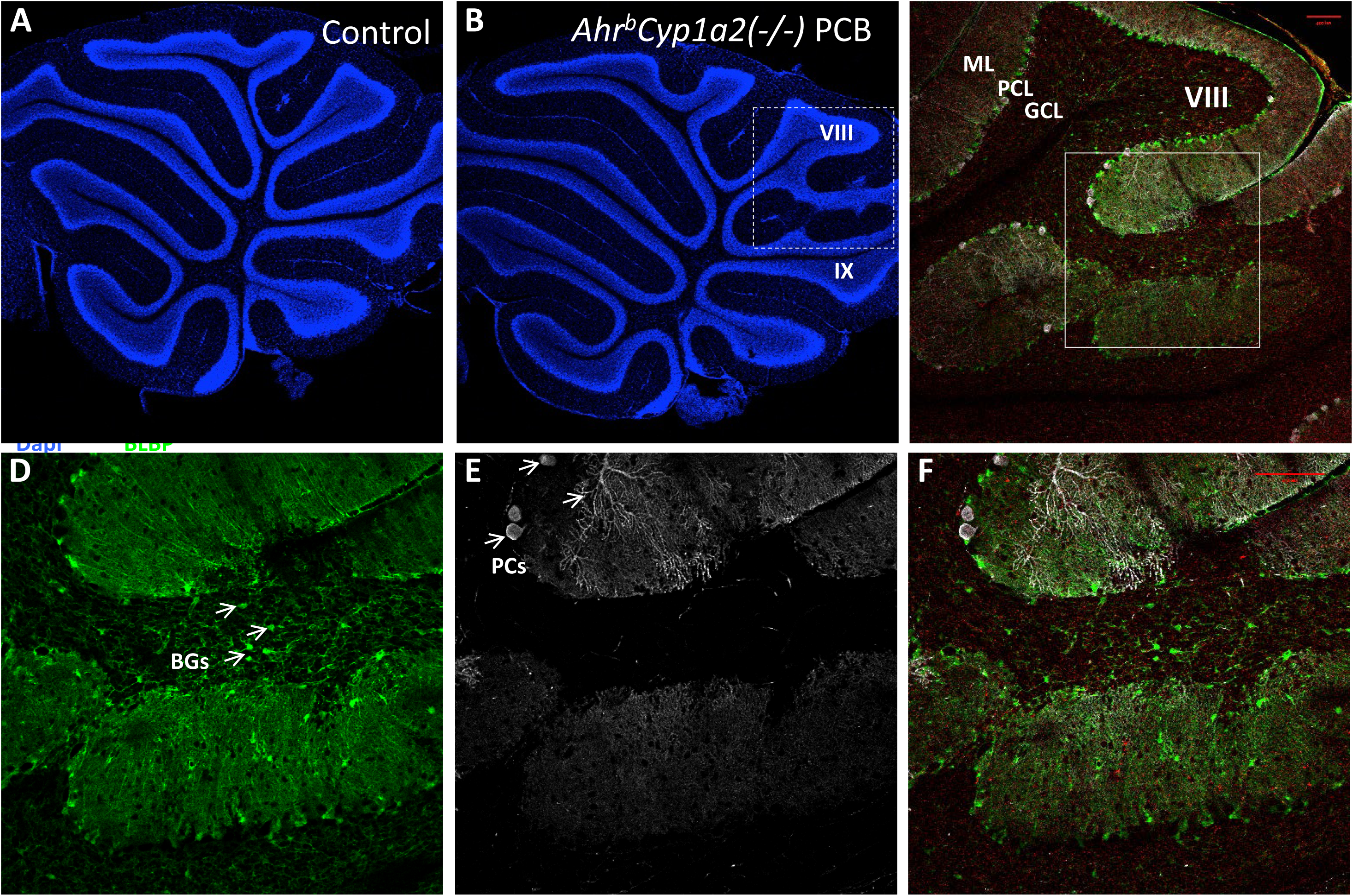
A-B, representative image of a knock out-control mouse (A) and knock out-treated with PCB (B) stained with DAPI. C, Zoom of the dashed area in B, showing the aberrant lamination between folias VIII and IX. Note the continuity of the GCL. D-F, Zoom of the square in C showing the absence of Chat+ Purkinje cells (PCs, white) and the invasion of BLBP+ Bergmann glial cells (BGs, green) in the folia VIII of treated KO mice. ML; molecular layer; PCL; Purkinje cell layer; GCL; granule cell layer. Scale bar 100 μm

### Gene expression differences in cerebellum and cortex

Based on the histology and immunohistochemistry results, we focused our gene expression studies on the cerebellum and the cortex, because both play a role in regulating motor function and motor memory. All changes in gene expression were compared with corn oil-treated control mice of the *Ahr*^*b*^*Cyp1a2(+/+)* genotype (i.e. the C57BL/6J strain). Details of fold-changes and P-values can be found in Tables 1-3.

**Table 1.**
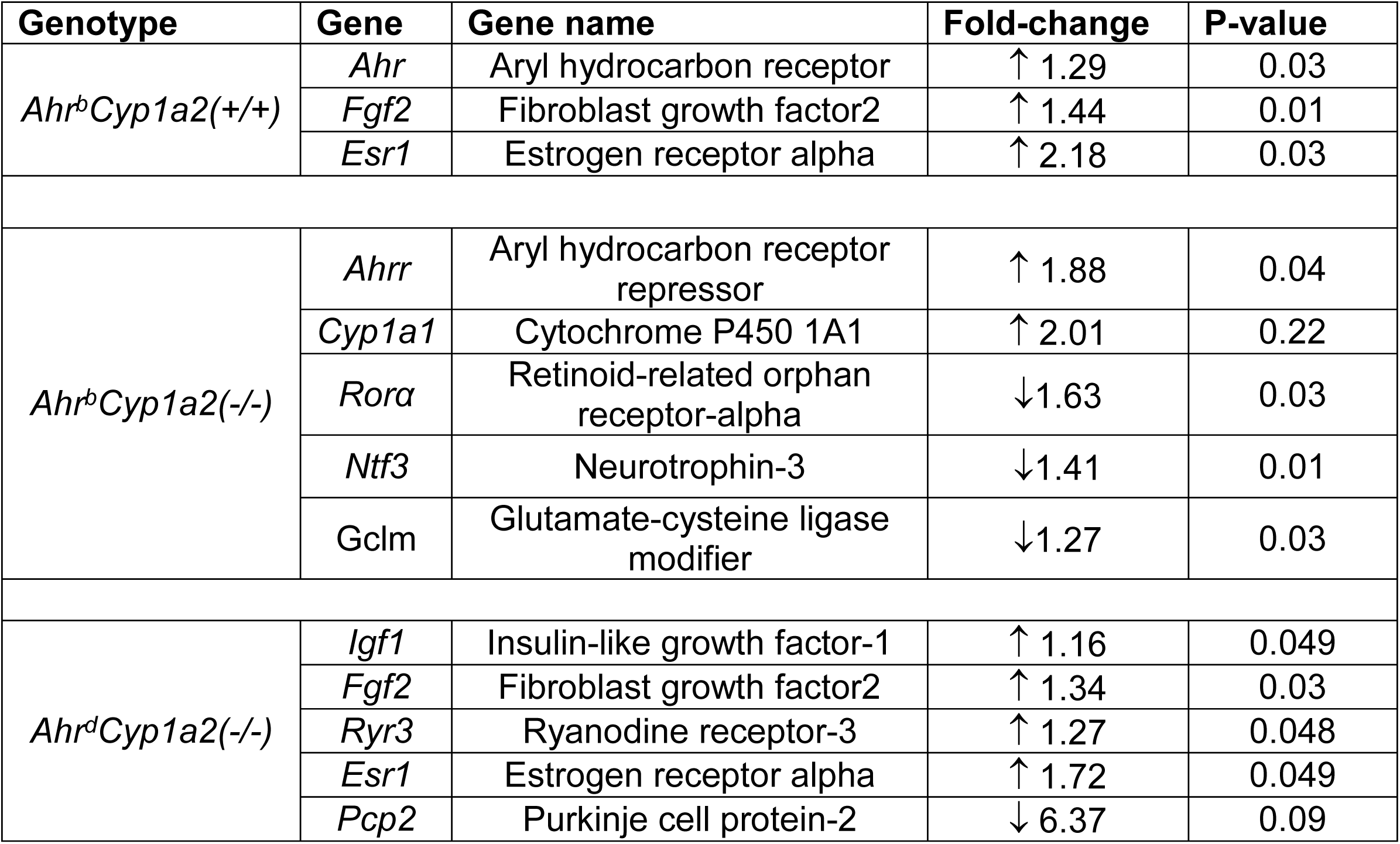
Gene expression differences in P14 cerebellum. Only genes with significant differences or fold-changes greater than 2 are shown. All values were compared against gene expression in corn oil-treated control B6 mice.

**Table 2.** Gene expression differences in P30 cerebellum. Only genes with significant differences or fold-changes greater than 2 are shown. All values were compared against gene expression in corn oil-treated control B6 mice.

|  |  |  |  |  |
| --- | --- | --- | --- | --- |
| *Ahr<sup>b</sup>Cyp1a2(+/+)* | *Cyp1b1* | Cytochrome P450 1B1 | ↑ 1.42 | 0.007 |
|  | *Grin2a* | NMDA receptor subtype 2A | ↑ 1.42 | 0.06 |
|  | *Ngf* | Nerve growth factor | ↑ 1.61 | 0.09 |
| *Ahr<sup>b</sup>Cyp1a2(-/-)* | *Ahr* | Aryl hydrocarbon receptor | ↓ 1.11 | 0.07 |
|  | *Cyp1a1* | Cytochrome P450 1A1 | ↑ 2.25 | 0.08 |
| *Ahr<sup>d</sup>Cyp1a2(-/-)* | *Ahr* | Aryl hydrocarbon receptor | ↓ 1.37 | 0.005 |
|  | *Cyp1a1* | Cytochrome P450 1A1 | ↑ 2.15 | 0.09 |
|  | *Pon2* | Paraoxonase-2 | ↓ 1.11 | 0.04 |

**Table 3.** Gene expression differences in P14 cortex. Only genes with significant differences or fold-changes greater than 2 are shown. All values were compared against gene expression in corn oil-treated control B6 mice.

|  |  |  |  |  |
| --- | --- | --- | --- | --- |
| *Ahr<sup>b</sup>Cyp1a2(+/+)* | *Ngf* | Nerve growth factor | ↓ 1.35 | 0.01 |
| *Ahr<sup>b</sup>Cyp1a2(-/-)* | *Cyp1a1* | Cytochrome P450 1A1 | ↑ 2.48 | 0.03 |
|  | *Cyp1b1* | Cytochrome P450 1B1 | ↑ 1.67 | 0.06 |
|  | *Mbp* | Myelin basic protein | ↓ 1.78 | 0.004 |
|  | *Grin2a* | NMDA receptor subtype 2A | ↓ 1.13 | 0.03 |
|  | *Ptgs2* | Prostaglandin G/H synthase-2 | ↓ 1.24 | 0.04 |
|  | *Pon2* | Paraoxonase-2 | ↓ 1.28 | 0.02 |
| *Ahr<sup>d</sup>Cyp1a2(-/-)* | *Ngf* | Nerve growth factor | ↓ 1.62 | 0.03 |
|  | *Esr1* | Estrogen receptor alpha | ↑ 1.30 | 0.049 |

### Differences at P14 in cerebellum

We found only three significant differences in PCB-treated *Ahr*^*b*^*Cyp1a2(+/+)*. Expression of the aryl hydrocarbon receptor and fibroblast growth factor-2 were modestly upregulated whereas expression of estrogen receptor alpha was more than doubled. Only *Ahr*^*b*^*Cyp1a2(-/-)* mice showed clear evidence of AHR activation with increases in both the aryl hydrocarbon receptor (1.88 fold increase) and CYP1A1 (2.01 increase), although the increase in CYP1A1 was not statistically significant. There was a significant down-regulation of retinoid-related orphan receptor-alpha (*Rorα*), neurotrophin-3 and glutamate-cysteine ligase modifier (*Gclm*). Gene expression in poor-affinity *Ahr*^*d*^*Cyp1a2(-/-)* mice had some similarities to wild type mice with significant increases in fibroblast growth factor-2 and estrogen receptor alpha. There were also significant increases in insulin-like growth factor-1 and ryanodine receptor-3. The greatest change was a more than 6-fold decrease in expression of Purkinje cell protein-2 (Pcp2); however the differences did not reach statistical significance (P = 0.09).

### Differences at P30 in cerebellum

Unexpectedly, all three lines of mice showed evidence of AHR activation in the cerebellum at P30, but the patterns of gene expression differed based on *Cyp1a2* genotype. CYP1B1 was significantly upregulated in *Ahr*^*b*^*Cyp1a2(+/+)* mice, whereas the AHR was down-regulated and CYP1A1 was upregulated in both *Ahr*^*b*^*Cyp1a2(-/-)* and *Ahr*^*d*^*Cyp1a2(-/-)* mice. We also found some evidence of compensatory processes with upregulation of both NMDA receptor subtype-2A (*Grin2a*) and nerve growth factor only in wild type *Ahr*^*b*^*Cyp1a2(+/+)* mice, but the differences did not reach statistical significance (P = 0.06 and P=0.09 respectively).

### Differences at P14 in cortex

The pattern of gene expression in cortex was much different. Nerve growth factor was significantly down-regulated in high-affinity *Ahr*^*b*^*Cyp1a2(+/+)* and poor-affinity *Ahr*^*d*^*Cyp1a2(-/-)* mice. Similar to P14, estrogen receptor alpha was also significantly upregulated in *Ahr*^*d*^*Cyp1a2(-/-)* mice. The most pronounced differences were in high-affinity *Ahr*^*b*^*Cyp1a2(-/-)* mice with clear evidence of AHR activation by significant upregulation of both CYP1A1 and CYP1B1. There was also significant down-regulation of myelin basic protein, Grin2a, prostagland G/H synthase-2 (*Ptgs2*) and paraoxonase-2 (*Pon2*).

### Tissue levels of PCB and hydroxylated metabolites

To determine if differential toxicokinetics could partially explain differences in gene expression, we compared levels of the parent PCB congeners in cerebellum, cortex, liver, adipose and blood (Table 4) and hydroxylated metabolites in blood and liver (Table 5). We found strikingly high levels of coplanar PCB 77 in adipose, liver, and blood of poor-affinity *Ahr*^*d*^*Cyp1a2(-/-)* mice and also significantly higher levels in cerebellum with no difference in cortex. Levels of coplanar PCB 126 were significantly higher in the liver of high-affinity *Ahr*^*b*^*Cyp1a2(+/+)* mice. Levels of coplanar PCB 169 were also higher in this line, although the differences didn’t reach statistical significance. In contrast, PCB 126 levels were significantly higher in the blood and adipose of the two *Cyp1a2(-/-)* knockout lines of mice.

**Table 4.** Tissue levels of PCBs in three genetically different pup groups. Values are presented as mean ± standard error in ng/g wet weight. * P < 0.05, ** P < 0.01, **A;* P < 0.001

| Tissue | Congener | *Ahr<sup>b</sup>Cyp1a2(+/+)* | *Ahr<sup>b</sup>Cyp1a2(-/-)* | *Ahr<sup>d</sup>Cyp1a2(-/-)* |
| --- | --- | --- | --- | --- |
| Adipose | PCB 77 | ND | 9 ± 2 | 14000 ± 2400*** |
|  | PCB 126 | 35 ± 5 | 95 ± 20** | 140 ± 34** |
|  | PCB 169 | 1200 ± 410 | 1700 ± 150 | 1600 ± 350 |
|  | PCB 118 | 17000 ± 4200 | 31000 ± 4400 | 33000 ± 6800 |
|  | PCB 105+153 | 30000 ± 3200 | 56000 ± 15000* | 77000 ± 19000* |
|  | PCB 138 | 22000 ± 4100 | 27000 ± 3100 | 26000 ± 5900 |
|  | PCB 180 | 25000 ± 6600 | 32000 ± 2400 | 30000 ± 6400 |
| Liver | PCB 77 | ND | ND | 730 ± 90*** |
|  | PCB 126 | 140 ± 40* | 13 ± 4 | 6 ± 1 |
|  | PCB 169 | 600 ± 300 | 150 ± 30 | 50 ± 7 |
|  | PCB 118 | 840 ± 280 | 1800 ± 220 | 980 ± 120 |
|  | PCB 105+153 | 1800 ± 350 | 3200 ± 310** | 3600 ± 400** |
|  | PCB 138 | 1400 ± 220 | 2200 ± 140* | 1300 ± 120 |
|  | PCB 180 | 1600 ± 420 | 3100 ± 200* | 1500 ± 180 |
| **Cerebellum** | PCB 77 | ND | ND | 160 ± 30*** |
|  | PCB 126 | ND | 13 ± 2 | ND |
|  | PCB 169 | 3 ± 1 | 10 ± 4 | 3 ± 1 |
|  | PCB 118 | 130 ± 20 | 300 ± 40* | 200 ± 30 |
|  | PCB 105+153 | 180 ± 40 | 380 ± 30* | 250 ± 40 |
|  | PCB 138 | 160 ± 30 | 270 ± 30* | 140 ± 20 |
|  | PCB 180 | 200 ± 40 | 370 ± 30* | 210 ± 30 |
| **Cortex** | PCB 77 | 100 ± 88 | 13 ± 2 | 140 ± 100 |
|  | PCB 118 | 110 ± 43 | 140 ± 70 | 190 ± 150 |
|  | PCB 126 | 3 ± 1 | ND | ND |
|  | PCB 169 | 9 ± 4 | 4 ± 2 | ND |
|  | PCB 105+153 | 150 ± 60 | 190 ± 90 | 260 ± 210 |
|  | PCB 138 | 150 ± 50 | 130 ± 60 | 140 ± 110 |
|  | PCB 180 | 170 ± 73 | 160 ± 70 | 220 ± 170 |
| **Blood** | PCB 77 | 33 ± 7 | ND | 92 ± 5*** |
|  | PCB 126 | ND | 2 ± 1 | ND |
|  | PCB 169 | 4 ± 1 | 7 ± 1*** | 2 ± 1 |
|  | PCB 118 | 46 ± 5 | 70 ± 5*** | 75 ± 4*** |
|  | PCB 105+153 | 104 ± 24 | 190 ± 10*** | 85 ± 5 |
|  | PCB 138 | 70 ± 4 | 74 ± 4 | 83 ± 5 |
|  | PCB 180 | 95 ± 9 | 110 ± 4* | 80 ± 6 |

**Table 5.** Blood and liver levels of OH-PCBs in three genetically different pup groups. Values are presented as mean ± standard error in ng/g wet weight.

| **Tissue** | **Congener** | ***Ahr<sup>b</sup>Cyp1a2*(+/+)** | ***Ahr<sup>b</sup>Cyp1a2*(-/-)** | ***Ahr<sup>d</sup>Cyp1a2*(-/-)** |
| --- | --- | --- | --- | --- |
| **Blood** | 5-77 + 4-107 | 170 ± 70** | 11 ± 1 | 2 ± 1 |
| | 4-146 | 4 ± 1 | 6 ± 1 | 2 ± 1 |
|  | 3-138 | ND | ND | ND |
| **Liver** | 5-77 + 4-107 | 190 ± 30** | 56 ± 22 | 100 ± 20 |
| | 4-146 | 16 ± 6 | 16 ± 3 | 14 ± 3 |
| | 3-138 | 15 ± 5 | 13 ± 4 | 9 ± 2 |

When comparing levels of noncoplanar PCB congeners, we found significantly higher levels of PCB105+PCB153 in the blood and cerebellum of *Ahr*^*b*^*Cyp1a2(-/-)* mice and significantly higher levels in the liver and adipose of both *Cyp1a2(-/-)* knockout lines. Levels of PCB 118 were significantly higher in the cerebellum of *Ahr*^*b*^*Cyp1a2(-/-)* mice and significantly higher in the blood of both *Cyp1a2(-/-)* knockout lines. Levels of PCB 138 were significantly higher in the cerebellum of *Ahr*^*b*^*Cyp1a2(-/-)* mice with no significant differences in other tissues. Levels of PCB 180 were significantly higher in the cerebellum, blood and liver of *Ahr*^*b*^*Cyp1a2(-/-)* mice with no differences in cortex and adipose.

## Discussion

Although the neurotoxicity of PCBs has been well established (Jacobson et al. 1997; Jacobson et al. 2003; Schantz et al. 2003; Ross 2004), less is known about the cause of motor deficits following PCB exposure, and there is minimal data concerning genetic susceptibility to PCBs. Of specific concern to human health is the possibility that PCBs may be a risk factor for Parkinson’s disease (Steenland et al. 2006; Caudle et al. 2008) and reports of motor deficits in children exposed to high levels during early brain development (Stewart et al. 2000; Vreugdenhil et al. 2002; Wilhelm et al. 2008). Because human body burdens are expected to remain high for several generations (Quinn et al. 2011), understanding genetic susceptibility will be important in identifying and protecting the most vulnerable human populations from PCB-induced motor dysfunction.

Our prior work looking at developmental PCB neurotoxicity in mice revealed that differences in both the aryl hydrocarbon receptor and CYP1A2 affect susceptibility to PCB-induced learning and memory deficits (Curran et al. 2011b, 2012). We have now extended those studies to PCB-induced motor dysfunction and found similar genetic differences in susceptibility (Stegman et al. 2014; Colter et al. 2017 submitted). The studies reported here were designed to identify the specific brain regions affected and the critical pathways involved in motor dysfunction. By assessing multiple endpoints, we were able to address three important questions: What is the evidence that PCBs are a risk factor for Parkinson’s disease? Can motor deficits be explained instead by thyroid hormone depletion and cerebellar dyfunction? Is there cross-talk between AHR-mediated pathways and other pathways involved in PCB neurotoxicity?

Immune suppression is a hallmark of AHR-mediated developmental toxicity, which can be easily assessed by measuring spleen and thymus weights in neonates (Lin et al. 2011; Curran et al. 2006, 2011a). We found strong evidence of immune suppression in the *Ahr*^*b*^*Cyp1a2(-/-)* mouse line previously identified as most susceptible to PCB-induced neurotoxicity (Figs. 2A-D). We also found significant upregulation of AHR target genes in this mouse line in both cerebellum and cortex (Tables 1-3) with more modest changes in AHR-regulated genes in the other lines of mice. Together, these findings support prior studies that maternal CYP1A2 is protective against AHR-mediated toxicity (Diliberto et al. 1997; Dragin et al. 2006; Curran et al. 2011a-b) and that high-affinity *Ahr*^*b*^ mice are more susceptible than poor-affinity *Ahr*^*d*^ mice (Curran et al. 2006, 2011a-b).

We compared thyroid hormone levels at P14 and at 4 months of age (Figs. 3A-C). Our findings were consistent with previous reports showing depletion of circulating thyroid hormone in neonatal mice (Banal and Zoeller 2008; Darras 2008; Curran et al. 2011b), but we found no difference in T4 levels in adult mice. The greatest depletion in T4 occurred in PCB-treated *Ahr*^*b*^*Cyp1a2(-/-)* mice, again suggesting this genotype is uniquely susceptible. In addition to previously described changes in AHR-regulated genes, the greatest changes seen in our qPCR arrays were in genes involved in thyroid hormone pathways. PCB-treated *Ahr*^*b*^*Cyp1a2(-/-)* mice had decreased expression of *Rorα,* which normally enhances thyroid hormone signaling (Qui et al. 2009) and is mutated in the staggerer mouse resulting abnormal cerebellar development (Hamilton et al. 1996). PCB-treated *Ahr*^*b*^*Cyp1a2(-/-)* mice also had significantly decreased levels of neurotrophin-3 at P14. Our findings are consistent with thyroid hormone depletion and those of Koibuchi et al. (2001) who reported decreased expression of Ntf-3 and Rorα in hypothyroid neonatal mice.

We also found differences in the expression of *Fgf2, Mbp, Ngf, and Pcp2*, which are all regulated, at least in part, by thyroid hormone (Koibuchi & Chin 2000). Interestingly, the direction of change varied across genotypes and brain regions. Upregulation of *Fgf2* and estrogen receptor-alpha in wild type *Ahr*^*b*^*Cyp1a2(+/+)* and poor-affinity *AhrdCyp1a2(-/-)* mice could indicate some compensatory mechanism in response to PCB exposure while the 6-fold down-regulation of *Pcp2* in *AhrdCyp1a2(-/-)* mice indicates a unique mode of action for PCB toxicity in that line.

Hydroxylated PCB metabolites appear most important in causing abnormal cerebellar development (Giera et al. 2011; Takekuchi et al. 2011; Berghuis et al. 2013), and there is evidence that CYP1A1 can metabolize non-coplanar PCBs (Gauger et al. 2007) altering the expression of thyroid hormone-regulated genes during human fetal development (Wadzinski et al. 2014). Koh et al. (2016) recently reported finding hydroxylated PCB metabolites in 60% of human sera samples, so we also examined PCB distribution and metabolism in weanling pups.

We found significant differences across genotypes with poor-affinity *AhrdCyp1a2(-/-)* mice showing strikingly high levels of coplanar PCB 77 in adipose, liver, and cerebellum. This was consistent with our previous findings in gavage-treated animals (Curran et al. 2011a). Wild-type *Ahr*^*b*^*Cyp1a2(+/+)* mice had significantly higher levels of coplanar PCB 126 in liver whereas both *Cyp1a2(-/-)* lines of mice had higher levels in adipose (Table 4). This is also consistent with our previous findings and evidence of coplanar PCB sequestration by maternal CYP1A2. However, there were no significant differences in the production of hydroxylated PCB metabolites (Table 5). Since our previous work demonstrated differences in parent congener levels throughout gestation and lactation, it will be important to extend these studies to other development time points and to assess both parent congeners and hydroxylated PCB metabolites.

Our histological analysis and immunohistochemistry experiments focused on effects in the cerebellum and striatum to determine which region was most affected by developmental PCB exposure. We found no differences in the area and perimeter of the cerebellum or granule cell layer and no difference in Purkinje cell numbers. However, we discovered a foliation defect between folia VIII and IX that was significantly more prevalent in PCB-treated *Cyp1a2(-/-)* mice (Figs. 6A-F). This was a novel finding for PCB-treated mice, although the defect has been reported in the background B6 strain previously (Van Dine et al. 2015). Still, Faquier et al. (2014) clearly identified Begmann glia and Purkinje cells as targets of thyroid hormone receptor-alpha during postnatal cerebellum development. Disruption is normal signaling resulted in abnormal granule cell migration and differentiation. In contrast, we found no differences in tyrosine hydroxylase staining in the striatum indicating no substantial changes in nigrostriatal dopaminergic pathways. Together, these data suggest that the primary target of PCB-induced motor dysfunction is the developing cerebellum and that the mechanism of toxicity is linked to disruptions in normal thyroid hormone signaling. In fact, motor dysfunction has been previously associated with defects in the development of folias VIII-IX of the cerebellum (López-Juárez et al. 2011).

We found no evidence of oxidative stress or neuro-inflammation at the time points tested and no evidence that genes important to calcium homeostasis were changed. There was a significant, but modest 1.27-fold increase in expression of ryanodine receptor-3 in the cerebellum of *Ahr*^*d*^*Cyp1a2(-/-)* mice at P14. The functional significance of that change is uncertain, but the findings are worth further exploration since the ryanodine receptor is strongly linked to PCB neurotoxicity (Pessah et al. 2010).

## Conclusions

We conclude that while all mouse lines experienced some level of PCB developmental toxicity, high-affinity *Ahr*^*b*^*Cyp1a2(-/-)* mice were most susceptible and *Ahr*^*b*^*Cyp1a2(+/+)* mice were most resistant. AHR activation appears to be a key difference between the two knockout lines with distinct gene expression patterns seen in *Ahr*^*b*^*Cyp1a2(-/-)* v. *Ahr*^*d*^*Cyp1a2(-/-)* mice and significant differences in PCB distribution and metabolism as well. We found multiple lines of evidence converging on the cerebellum as the primary target organ for PCB-induced motor deficits with disruption of normal thyroid hormone signaling the most likely mode of action. There was minimal evidence that developmental PCB exposure is a risk factor for Parkinson’s disease. However, this does not rule out the possibility of neurotoxicity in adults exposed over a longer period of time or to higher doses from occupational exposures.

## Acknowledgements

This work was supported by the National Institutes of Health (R15ES020053, P20 GM103436, ES05605 and ES013661), the National Science Foundation (RSF-034-07, DUE-STEP-096928), and the following grants from Northern Kentucky University: Faculty Development Project Grants, College of Arts & Sciences Collaborative Faculty-Student Award, Center for Integrative Natural Science and Mathematics Research Grants, and Dorothy Westerman Herrmann funds. We thank Joshua Lambert, University of Kentucky Department of Statistics, Jiaying Weng and Ya Qi, University of Kentucky Department of Statistics Applied Statistics Laboratory, for assistance with data analysis, and we acknowledge the generous donation of *Cyp1a2(-/-)* knockout mice from Dr. Daniel W. Nebert, University of Cincinnati Department of Environmental Health.

## Data availability

The data sets for all figures and tables in this article are available from the corresponding author on reasonable request.

## Animal welfare

All applicable international, national, and/or institutional guidelines for the care and use of animals were followed, and all protocols were approved by the Northern Kentucky University Institutional Animal Care and Use Committee.

---

**Table S1.** Environmentally relevant mixture of PCB congeners used to treat pregnant mice.

| **Congener** | **Chemical Formula** | **Planarity** | **Dose (per kg/day)** |
| --- | --- | --- | --- |
| PCB 77 | 3,3',4,4'-Tetrachlorobiphenyl | Coplanar | 0.5 mg |
| PCB 126 | 3,3',4,4',5-Pentachlorobiphenyl | Coplanar | 2.5 µg |
| PCB 169 | 3,3',4,4',5,5'-Hexachlorobiphenyl | Coplanar | 25 µg |
| PCB 105 | 2,3,3',4,4'-Pentachlorobiphenyl | Non-coplanar | 1 mg |
| PCB 118 | 2,3',4,4',5-Pentachlorobiphenyl | Non-coplanar | 1 mg |
| PCB 138 | 2,2',3,4,4',5'-Hexachlorobiphenyl | Non-coplanar | 1 mg |
| PCB 153 | 2,2',4,4',5,5'-Hexachlorobiphenyl | Non-coplanar | 1 mg |
| PCB 180 | 2,2',3,4,4',5,5'-Heptachlorobiphenyl | Non-coplanar | 1 mg |

**Table S2.**
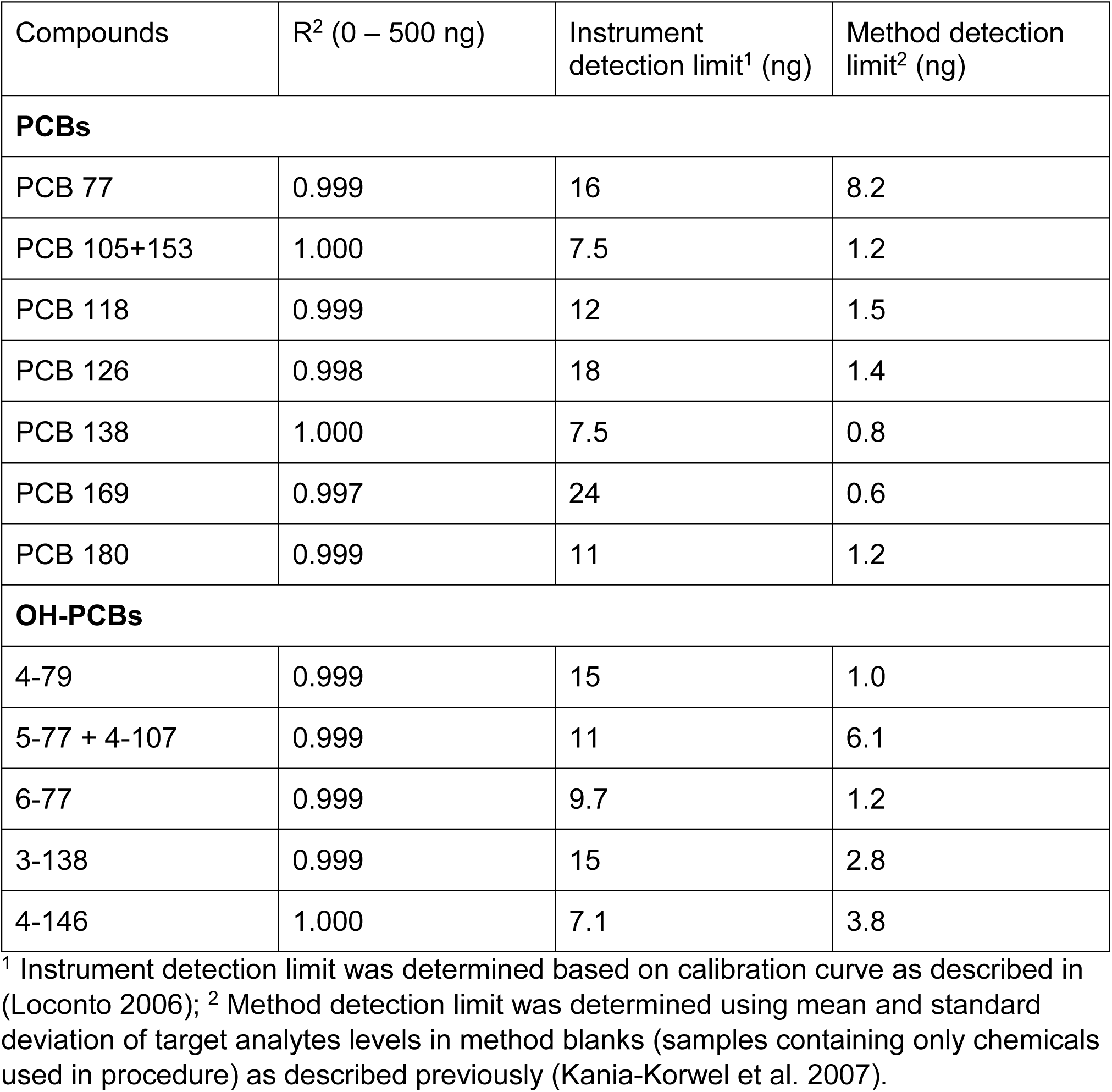
Linear range and R^2^, instrument and method detection limit for all target analytes in this study.

**Table S3.** Background PCB and OH-PCB levels (ng/g wet weight) in animals exposed to vehicle alone (corn oil).

| Compounds | Adipose<br>(n=3) | Blood<br>(n=8) | Cerebellum<br>(n=3) | Cortex<br>(n=3) | Liver<br>(n=3) |
| --- | --- | --- | --- | --- | --- |
| **PCBs** |  |  |  |  |  |
| PCB 77 | ND | 8.5 ± 5.9 | 34 ± 12 | 0.57 ± 0.74 | 4.7 ± 1.0 |
| PCB 105+153 | 163 ± 80 | 2.4 ± 2.0 | 1.7 ± 1.9 | 0.40 ± 0.18 | 68 ± 93 |
| PCB 118 | 107 ± 53 | 0.66 ± 0.36 | 6.0 ± 3.0 | 0.59 ± 0.73 | 5.6 ± 4.3 |
| PCB 126 | ND | 0.37 ± 0.42 | 5.0 ± 7.3 | 1.3 ± 1.6 | 2.0 ± 0.7 |
| PCB 138 | 54 ± 14 | 1.6 ± 2.9 | 4.1 ± 0.5 | 0.48 ± 0.50 | 7.5 ± 4.4 |
| PCB 169 | 6.6 | 0.52 ± 0.67 | 1.2 ± 1.4 | 0.58 ± 0.71 | 3.6 ± 6.0 |
| PCB 180 | 53 ± 16 | 1.3 ± 1.5 | 4.1 ± 0.6 | 0.16 ± 0.10 | 8.1 ± 3.3 |
| **OH-PCBs** |  |  |  |  |  |
| 4-79 | ND | 0.21 ± 0.08 | 2.2 ± 0.9 | ND | 0.7 ± 0.9 |
| 5-77 + 4-107 | 21 ± 6 | 1.8 ± 1.4 | 16 ± 4 | 3.4 ± 4.3 | 6.3 ± 2.5 |
| 6-77 | ND | 0.71 ± 0.36 | 9.6 ± 7.4 | ND | 0.22 ± 0.01 |
| 3-138 | ND | 0.21 ± 0.10 | 1.4 ± 1.6 | 0.74 ± 1.2 | 0.30 ± 0.24 |
| 4-146 | ND | 0.14 ± 0.08 | 7.2 ± 8.1 | 1.4 ± 2.2 | 0.08 ± 0.08 |

ND – not detected.

## References

[ATSDR] Agency for Toxic Substances and Disease Registry. (2015). 2015 CERCLA National Priorities List of Hazardous Substances. U.S. Department of Health and Human Services, Public Health Service. Atlanta, GA. https://www.atsdr.cdc.gov/spl/

Caudle, W. M., Richardson, J. R., Delea, K. C., Guillot, T. S., Wang, M., Pennell, K. D. Miller GW. (2006). Polychlorinated biphenyl-induced reduction of dopamine transporter expression as a precursor to Parkinson’s disease-associated dopamine toxicity. Toxicol.Sci., 92, 490–499.

Curran, C. P., Miller, K. A., Dalton, T. P., Vorhees, C. V., Miller, M. L., Shertzer, H. G. et al. (2006). Genetic differences in lethality of newborn mice treated in utero with coplanar versus non-coplanar hexabromobiphenyl. Toxicol.Sci., 89, 454–464.

Curran, C.P., Vorhees, C.V., Williams, M.T., Genter, M.B., Nebert, D.W. (2011a). In utero and lactational exposure to a complex mixture of polychlorinated biphenyls: Toxicity in pups dependent on the Cyp1a2 and Ahr genotypes. Toxicol. Sci. 119, 189–208.

Curran, C.P. Nebert D.W., Genter M.B., Patel K.V., Schafer, T.L., Skelton M.R., Williams M.T. and Vorhees C.V. (2011b). In utero and lactational exposure to PCBs in mice: Adult offspring show altered learning and memory depending on *Cyp1a2* and *Ahr* genotypes. Enironv. Health Perspect. 119(9), 1286–93.

Curran C.P., Altenhofen E.,. Ashworth A.A.,. Brown A., Curran M.A., Evans A., Floyd R., Fowler J.P., Garber H., Hays B., Kamau-Cheggeh C., Kraemer S., Lang A.L., Mynhier A., Samuels A. and Strohamier C. (2012). *AhrdCyp1a2(-/-)* mice show increased susceptibility to PCB-induced developmental neurotoxicity. Neurotoxicology. 33(6), 1436–42.

Darras V.M. (2008). Endocrine disrupting polyhalogenated organic pollutants interfere with thyroid hormone signalling in the developing brain. The Cerebellum. 26-37.

Diliberto, J. J., Burgin, D., & Birnbaum, L. S. (1997). Role of CYP1A2 in hepatic sequestration of dioxin: studies using CYP1A2 knock-out mice. Biochem. Biophys.Res.Commun., 236, 431–433.

Dong, H., Wade, M., Williams, A., Lee, A., Douglas, G. R., & Yauk, C. (2005). Molecular insight into the effects of hypothyroidism on the developing cerebellum. Biochem. Biophys.Res.Commun., 330, 1182–1193.

Dragin, N., Dalton, T. P., Miller, M. L., Shertzer, H. G., & Nebert, D. W. (2006). For dioxin-induced birth defects, mouse or human CYP1A2 in maternal liver protects whereas mouse CYP1A1 and CYP1B1 are inconsequential. J.Biol.Chem., 281, 18591–18600.

Fauquier, T., Chatonnet F., Picou, F., Richard S., Fossat N., Aguilera N., Lamonerie T., and Flamant F., (2014). Purkinje cells and Bergmann glia are primary targets of the TRα1 thyroid hormone receptor during mouse cerebellum postnatal development. Development. 141, 166–175.

Gauger, K. J., Giera, S., Sharlin, D. S., Bansal, R., Iannacone, E., & Zoeller, R. T. (2007). Polychlorinated biphenyls 105 and 118 form thyroid hormone receptor agonists after cytochrome P4501A1 activation in rat pituitary GH3 cells. Environ.Health Perspect., 115, 1623–1630.

Giera S, Bansal R, Ortiz-Toro TM, Taub DG, Zoeller RT,. (2011). Individual polychlorinated biphenyl (PCB) congeners produce tissue- and gene-specific effects on thyroid hormone signaling during development. Endocrinology. 152(7):2909–19.

Goldey ES and Crofton KM. (1998). Thyroxine replacement attenuates hypothyroxinemia, hearing loss, and motor deficits following developmental exposure to Aroclor 1254 in rats. Toxicol Sci. 1998 Sep;45(1):94–105.

Gomara, B., Bordajandi, L. R., Fernandez, M. A., Herrero, L., Abad, E., Abalos, M., Rivera, J, González, MJ. (2005). Levels and trends of polychlorinated dibenzo-p-dioxins/furans (PCDD/Fs) and dioxin-like polychlorinated biphenyls (PCBs) in Spanish commercial fish and shellfish products, 1995-2003. J.Agric.Food Chem., 53, 8406–8413.

González-Franco DA, Ramírez-Amaya V, Joseph-Bravo P, Prado-Alcalá RA, and Quirarte GL. (2017). Differential Arc protein expression in dorsal and ventral striatum after moderate and intense inhibitory avoidance training. Neurobiology of Learning and Memory. 40:17– 26.

Hamilton BA, Frankel WN, Kerrebrock AW, Hawkins TL, FitzHugh W, Kusumi K, Russell LB, Mueller KL, van Berkel V, Birren BW, Kruglyak L, Lander ES. (1996). Disruption of the nuclear hormone receptor RORα in staggerer mice. Nature 379:736–739.

Hu D, Hornbuckle KC. (2010). Inadvertent polychlorinated biphenyls in commercial paint pigments. Environ Sci Technol. 44(8):2822–7.

Jacobson, J. L. & Jacobson, S. W. (1997). Evidence for PCBs as neurodevelopmental toxicants in humans. Neurotoxicology, 18, 415–424.

Jacobson, J. L. & Jacobson, S. W. (2003). Prenatal exposure to polychlorinated biphenyls and attention at school age. J.Pediatr., 143, 780–788.

Kania-Korwel I, Parkin S, Robertson LW, Lehmler H-J (2004) Synthesis of polychlorinated biphenyls and their metabolites with a modified Suzuki-coupling. Chemosphere 56: 735– 744

Kania-Korwel I, Hornbuckle KC, Peck A, Ludewig G, Robertson LW, Sulkowski WW, Espandiari P, Gairola CG, Lehmler H-J (2005) Congener specific tissue distribution of Aroclor 1254 and a highly chlorinated environmental PCB mixture in rats. Environ. Sci. Technol. 39: 3513–3520

Kania-Korwel I, Shaikh N, Hornbuckle KC, Robertson LW, Lehmler H-J (2007) Enantioselective disposition of PCB 136 (2,2’,3,3’,6,6’-hexachlorobiphenyl) in C57BL/6 mice after oral and intraperitoneal administration. Chirality 19: 56–66

Kania-Korwel I, Zhao H, Norstrom K, Li X, Hornbuckle, KC Lehmler HJ (2008) Simultaneous extraction and clean-up of polychlorinated biphenyls and their metabolites from small tissue samples using pressurized liquid extraction. J. Chromatogr. A 1214: 37–46

Koh, WX Hornbuckle, KC Marek, RF Wang, K Thorne, PS. (2016). Hydroxylated polychlorinated biphenyls in human sera from adolescents and their mothers living in twoU.S. Midwestern communities. Chemosphere. 147:389–95.

Koibuchi N, Chin WW. (2000). Thyroid hormone action and brain development. Trends Endocrinol Metab. 11(4):123–8.

Langer, P., Kocan, A., Tajtakova, M., Petrik, J., Chovancova, J., Drobna, B. et al. (2007). Fish from industrially polluted freshwater as the main source of organochlorinated pollutants and increased frequency of thyroid disorders and dysglycemia. Chemosphere, 67, S379– S385.

Lehmler H-J, Robertson LW (2001) Synthesis of polychlorinated biphenyls (PCBs) using the Suzuki-coupling. Chemosphere 45: 137–143

Lin P, Hu SW, Chang TH. (2003) Correlation between gene expression of aryl hydrocarbon receptor (AhR), hydrocarbon receptor nuclear translocator (*Arnt*), cytochromes P4501A1 (CYP1A1) and 1B1 (CYP1B1), and inducibility of CYP1A1 and CYP1B1 in human lymphocytes. Toxicol Sci.71(1):20–6.

Lin TM, Ko K, Moore RW, Buchanan DL, Cooke PS, Peterson RE. (2001). Role of the aryl hydrocarbon receptor in the development of control and 2,3,7,8-tetrachlorodibenzo-p-dioxin-exposed male mice. J Toxicol Environ Health A. 64(4):327–42.

Loconto PR (2006) Trace Environmental Quantitative Analysis. CRC Press, Boca Raton, FL.

López-Juárez A, Morales-Lázaro S, Sánchez-Sánchez R, Sunkara M, Lomelí H, Velasco I, Morris AJ, Escalante-Alcalde D. (2011). Expression of LPP3 in Bergmann glia is required for proper cerebellar sphingosine-1-phosphate metabolism/signaling and development. Glia. 59(4):577–89.

Maervoet J, Covaci A, Schepens P, Sandau CD, Letcher R (2004) A reassessment of the nomenclature of polychlorinated biphenyl (PCB) metabolites. Environ. Health Perspect. 112: 291–294

Marek RF, Thorne PS, Herkert NJ, Awad AM, Hornbuckle KC. (2017). Airborne PCBs and OH-PCBs inside and outside urban and rural U.S. schools. Environ Sci Technol. 51(14):7853– 7860.

Nebert, D. W. & Dalton, T. P. (2006). The role of cytochrome P450 enzymes in endogenous signalling pathways and environmental carcinogenesis. Nat.Rev.Cancer, 6, 947–960.

Qiu CH, Miyazaki W, Iwasaki T, Londoño M, Ibhazehiebo K, Shimokawa N, Koibuchi N. (2009). Retinoic Acid receptor-related orphan receptor alpha-enhanced thyroid hormone receptor-mediated transcription requires its ligand binding domain which is not, by itself, sufficient: possible direct interaction of two receptors. Thyroid. 19(8):893–8.

Quinn, C. L., Wania, F., Czub, G., & Breivik, K. (2011). Investigating intergenerational differences in human PCB exposure due to variable emissions and reproductive behaviors. Environ.Health Perspect., 119, 641–646.

Pessah, I. N., Cherednichenko, G., & Lein, P. J. (2010). Minding the calcium store: Ryanodine receptor activation as a convergent mechanism of PCB toxicity. Pharmacol. Ther., 125, 260–285.

Petersen, M. S., Halling, J., Bech, S., Wermuth, L., Weihe, P., Nielsen, F. et al. (2008). Impact of dietary exposure to food contaminants on the risk of Parkinson’s disease. Neurotoxicology, 29, 584–590.

Schantz, S. L. Widholm, J. J. & Rice, D. C. (2003). Effects of PCB exposure on neuropsychological function in children. Environ.Health Perspect., 111, 357–576.

Schramm H, Robertson LW, Oesch F (1985) Differential regulation of hepatic glutathione transferase and glutathione peroxidase activities in the rat. Biochem. Pharmacol. 34: 3735–3739.

Seegal, R. F., Brosch, K. O., & Bush, B. (1986). Polychlorinated biphenyls produce regional alterations of dopamine metabolism in rat brain. Toxicol.Lett., 30, 197–202.

Seegal, R. F., Brosch, K. O., & Okoniewski, R. J. (1997). Effects of in utero and lactational exposure of the laboratory rat to 2,4,2′,4′- and 3,4,3′,4′-tetrachlorobiphenyl on dopamine function. Toxicol.Appl.Pharmacol., 146, 95–103.

Seegal, R. F., Brosch, K. O., & Okoniewski, R. J. (2005). Coplanar PCB congeners increase uterine weight and frontal cortical dopamine in the developing rat: implications for developmental neurotoxicity. Toxicol.Sci., 86, 125–131.

Seegal, R. F., Bush, B., & Brosch, K. O. (1994). Decreases in dopamine concentrations in adult, non-human primate brain persist following removal from polychlorinated biphenyls. Toxicology, 86, 71–87.

Steenland, K., Hein, M. J., Cassinelli, R. T., Prince, M. M., Nilsen, N. B., Whelan, E. A. et al. (2006). Polychlorinated biphenyls and neurodegenerative disease mortality in an occupational cohort. Epidemiology, 17, 8–13.

Stegman M., Curran, C.P., Infante, S.K., Kromme, M., Hays, B., Taylor, K., Garber, H., and Lang, A. (2014) Assessing genetic susceptibility to motor function deficits following developmental exposure to polychlorinated biphenyls. Toxicol. Sci. Supp. 138(1): 457– 458.

Takahashi M., Negishi T. and Tashior T. (2008). Identification of genes mediating thyroid hormone action in the developing mouse cerebellum. J. of Neurochem. 104, 640–652.

Takeuchi S., Shiraishi F., Kitamura S., Kuroki H., Jin K., and Kojima H. (2011). Characterization of steroid hormone receptor activities in 100 hydroxylated polychlorinated biphenyls, including congeners identified in humans. Toxicology. 289, 112–121.

van den Hurk P, Kubiczak GA, Lehmler HJ, James MO (2002) Hydroxylated polychlorinated biphenyls as inhibitors of the sulfation and glucuronidation of 3-hydroxy-benzo[a]pyrene. Environ Health Perspect 110: 343–8.

Van Dine SE, Siu NY, Toia A, Cuoco JA, Betz AJ, Bolivar VJ, Torres G, Ramos RL. (2015). Spontaneous malformations of the cerebellar vermis: Prevalence, inheritance, and relationship to lobule/fissure organization in the C57BL/6 lineage. Neuroscience. 310:242–51.

Vreugdenhil, H. J., Lanting, C. I., Mulder, P. G., Boersma, E. R., & Weisglas-Kuperus, N. (2002). Effects of prenatal PCB and dioxin background exposure on cognitive and motor abilities in Dutch children at school age. J.Pediatr., 140, 48–56.

Zoeller R.T., and Rovett J. (2004). Timing of thyroid hormone action in the developing brain: Clinical observations and experimental findings. J. of Neuroendo. 16, 809–818.

